# Vimentin filaments integrate low complexity domains in a highly complex helical structure

**DOI:** 10.1101/2023.05.22.541714

**Authors:** Matthias Eibauer, Miriam S. Weber, Rafael Kronenberg-Tenga, Charlie T. Beales, Rajaa Boujemaa-Paterski, Yagmur Turgay, Suganya Sivagurunathan, Julia Kraxner, Sarah Köster, Robert D. Goldman, Ohad Medalia

## Abstract

Intermediate filaments (IFs) are integral components of the cytoskeleton. They provide cells with tissue-specific mechanical properties and are involved in numerous cellular processes. Due to their intricate architecture, a 3D structure of IFs has remained elusive. Here we use cryo-focused ion beam milling, cryo-electron microscopy and tomography, to obtain a 3D structure of vimentin IFs (VIFs). VIFs assemble into a modular, densely-packed and highly-ordered helical symmetric structure of 40 α-helices in cross-section, organized into 5 protofibrils. Surprisingly, the intrinsically disordered head domains form an amyloid-like fiber in the center of VIFs, while the intrinsically disordered tails form lateral connections between the protofibrils. Our findings demonstrate how protein domains of low sequence complexity can complement well-folded protein domains to construct a biopolymer with striking strength and stretchability.

## Introduction

Intermediate filaments (IFs) are expressed in a cell-type specific manner, providing the cytoskeleton with cell-type specific viscoelastic properties [1]. In humans, IF proteins are encoded by 70 genes [2], including lamins [3], keratins [4], and neuro-filaments [5], and are associated with a wide range of functions [6, 7] and at least 72 human pathologies [2, 8]. Vimentin is mainly expressed in cells of mesenchymal origin where it assembles into extensive, dynamic and hyperelastic filament networks [9, 10]. Individual vimentin IFs (VIFs) can be stretched up to 3.5 times of their initial length [11, 12], tolerating much higher strains and exhibiting greater flexibility than F-actin and microtubules [13]. VIFs are involved in various cytoskeletal processes, such as maintenance of cell shape [14, 15], development of focal adhesions [16], protrusion of lamellipodia [17], assembly of stress-fibres [18, 19], and signal transduction [20]. Vimentin expression and assembly of VIFs are markers and regulators of the epithelial-mesenchymal transition [21]. They are involved in cancer initiation, progression and metastasis [22-24], and several other pathophysiological conditions [25-28].

At the molecular level, vimentin monomers interact to form in-register, parallel dimers [29], ∼48 nm in length. Like all IF proteins [30], the dimers contain a central coiled-coil rod domain composed of consecutive α-helical segments, connected by flexible linkers. The rod domain is flanked by intrinsically disordered N-terminal head and C-terminal tail domains [31]. The early steps in VIF assembly involve the formation of elongated tetramers consisting of two anti-parallel, staggered dimers [32-34], ∼62 nm in length. The vimentin tetramer is the basic building block for subsequent unit-length filament formation and VIF assembly [35-37].

While the tetramer is well characterized, solving the 3D structure of fully assembled VIFs and IFs in general with conventional methodologies has proven to be a major challenge. This is likely attributable to the unique elongated shape of their tetrameric building blocks, which contain a substantial fraction of intrinsically disordered domains and flexible linkers. IFs also exhibit extensive structural polymorphism [38-40]. Therefore, the 3D structure of a fully assembled IF has remained elusive [41].

Here we utilized cryo-focused ion beam (cryo-FIB) milling followed by cryo-electron tomography (cryo-ET) to study VIFs in-situ. With this approach, we revealed their native polymerization state and measured their previously unknown helical symmetry parameters. We used this information to obtain 3D structures of VIFs assembled from full-length and tailless vimentin proteins, using single particle cryo-electron microscopy (cryo-EM). Based on these structures we have constructed a complete atomic model of VIFs. This model shows an unique molecular architecture comprising a highly-ordered helical symmetric scaffold surrounding a central amyloid-like fiber.

### Polymerization state in-situ

Mouse embryonic fibroblasts (MEFs) are widely used for investigating the cellular organization and function of VIFs (**Fig. 1A**). To study VIFs in their native cellular environment, we cultured MEFs on electron microscopy grids, vitrified the cells without chemical fixation, prepared thin sections of the MEFs by cryo-FIB milling [42-44], and imaged the resulting lamellas by cryo-ET (**Fig. 1B, Extended Data Table 1, and Supplementary Video 1**). The unique advantage of this approach is that it retains the molecular organization of the sample completely intact [45]. Missing wedge induced resolution anisotropy in the tomograms was reduced with IsoNet (**Extended Data Fig. 1A**) [46].

**Figure 1.**
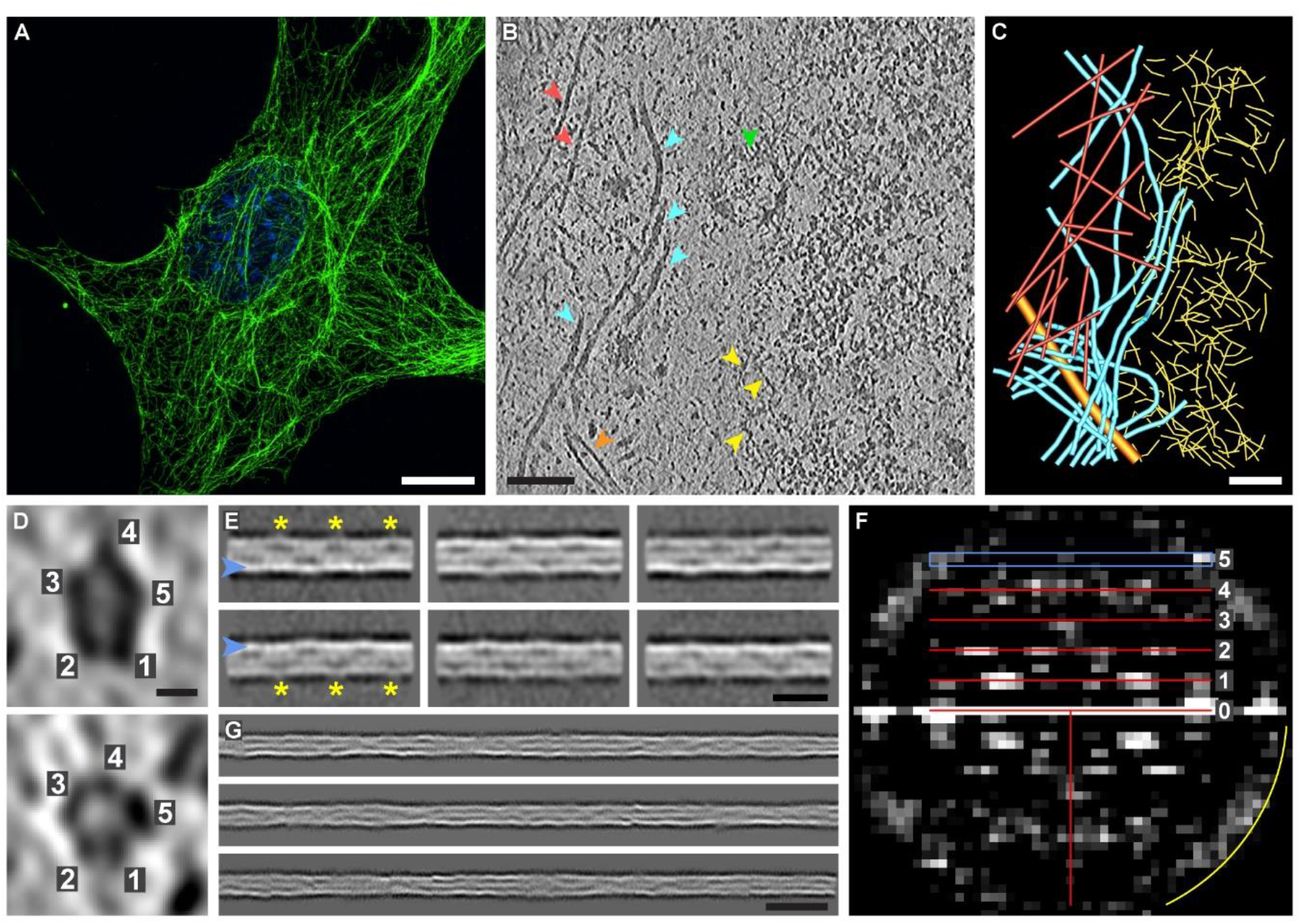
VIFs are built from 5 protofibrils in cells. (**A**) Slice through a 3D-SIM image of a MEF, fixed and stained with anti-vimentin (green) and the nucleus is stained with DAPI (blue). The VIF network extends over the whole cellular volume, with regions of lower and higher network density, and forms a cage around the nucleus. Scale bar is 10 µm. (**B**) Slice (8.84 Å thick) through a tomogram of a cryo-FIB milled MEF, recorded in a region around the nuclear envelope. VIFs are marked with cyan arrowheads. Actin filaments are labelled with red and lamin filaments with yellow arrowheads. Microtubule (MT), orange arrowhead; nuclear pore complex (NPC), green arrowhead. Scale bar is 100 nm. (**C**) Segmentation of the biopolymers present in (**B**). Scale bar is 100 nm. (**D**) Cross-sections of VIFs reveal 5 protofibrils. Scale bar is 5 nm. (**E**) Class averages of VIFs show a helical pattern with a repeat distance of ∼180 Å (yellow asterisks) and one side of the filament boundary appears pronounced in projection (blue arrowheads). Scale bar is 180 Å. (**F**) Combined power spectrum of class averages as shown in (**E**). The presence of layer lines confirms the helical architecture of VIFs. The first layer line appears at ∼1/185 Å and reflects the repeating pattern observed in the class averages. The layer line and peak distribution with a meridional reflection on the fifth layer line at ∼1/37 Å is compatible with a helical assembly of five subunits per repeat. The yellow arc in the power spectrum indicates 1/26 Å. (**G**) Gallery of three computationally assembled VIFs. This technique allows to follow the progression of VIFs with improved signal-to-noise ratio over a substantial length. Scale bar is 35 nm.

In regions around the nuclear envelope, the resulting tomograms show the complete set of filamentous biopolymers present in MEFs (**Fig. 1C**), including F-actin, microtubules and nuclear lamins [3]. VIFs can be clearly distinguished based on their tubular shape with a diameter of ∼11 nm. Contrary to expectations based on previous work [47], VIF cross-sections suggest that they are built from 5 protofibrils (**Fig. 1D and Extended Data Fig. 1B**).

### Determination of helical symmetry

In order to verify the pentameric cross-sections of VIFs, MEFs were cultured on electron microscopy grids as described before, but this time treated with a permeabilization procedure that preserves the VIF network (**Extended Data Fig. 2A**), while depleting soluble components from the cytoplasm and the nucleus [3]. Subsequently, the detergent-treated MEFs were vitrified and imaged by cryo-ET (**Extended Data Fig. 2B, Extended Data Table 2, and Supplementary Video 2**). This sample preparation results in thinned MEFs and therefore VIFs from different z-heights are concentrated in the tomograms. This procedure allows for an extensive statistical analysis of the filaments in a fraction of the time required with the cryo-FIB approach.

In total, we extracted ∼390,000 overlapping segments of VIFs with a length of 65 nm from the tomograms and subjected this data to a 2D classification procedure [48, 49]. The class averages reveal a clear periodic pattern (**Fig. 1E, yellow asterisks**) with a repeat distance of ∼180 Å (**Extended Data Fig. 2C**), as measured by autocorrelation analysis [50]. Additionally, the class averages show that one filament wall appears more pronounced in projection than its counterpart (**Fig. 1E, blue arrowheads**), further supporting that an odd number of protofibrils builds the VIFs (**Extended Data Fig. 2D**).

Subsequently, we combined the periodic class averages into a power spectrum, which shows an apparent layer line pattern (**Fig. 1F**). This is a direct prove that VIFs are built with helical symmetry [50]. The first layer line is positioned at ∼1/185 Å, which corroborates the previous autocorrelation measurement (**Extended Data Fig. 2C**). Moreover, the Bessel peak distribution along the layer line spectrum is compatible with a helical assembly of 5 building blocks in one helical pitch. Consequently, on the fifth layer line, positioned at ∼1/37 Å (**Fig. 1F, blue rectangle**), a meridional peak can be detected. Therefore, we conclude that the unique pentameric cross-sections of VIFs as observed in the in-situ cryo-FIB/cryo-ET data are formed by the specific helical symmetry of VIFs, as independently verified with cryo-ET data obtained from detergent-treated MEF preparations.

### Computational filament assembly

To improve the precision of the helical symmetry measurement and allow for higher resolution analysis, we computationally assembled extended stretches of VIFs (**Fig. 1G, Extended Data Fig. 3**) [4]. Combining these long-range VIF observations into a single power spectrum improved the resolution of the layer lines significantly. In fact, the previously observed layer lines (**Fig. 1F**) decompose at high resolution into bundles of fine layer lines (**Extended Data Fig. 4C**). Comprehensive power spectrum analysis yielded a helical rise of 42.5 Å and a helical twist of 73.7° (**Extended Data Fig. 4**).

This analysis also indicates that VIFs are assembled from 4.88 tetramers, packed into a helical pitch length of 207.4 Å. This result recapitulates a characteristic feature of IFs, namely their axial periodicity of ∼21 nm observed through rotary shadowing techniques [1, 30, 51-53].

Since the mass-per-length of a helical filament is proportional to its helical rise [54], the observed VIFs exhibit a mass-per-length of 53.7 kDa/nm, in agreement with data obtained by scanning transmission electron microscopy [38, 39].

### Structure of VIFs

We then aimed to solve the 3D structure of fully assembled VIFs. To this end we polymerized VIFs in-vitro from bacterially expressed and purified human vimentin, and imaged the filaments with cryo-EM. We extracted ∼1.5 million VIF segments from the dataset, which were subjected to extensive 2D and 3D classifications (**Extended Data Fig. 5, A and B**) [55, 56]. These calculations independently converged to the identical (up to two decimal places) helical parameter set as measured by power spectrum analysis of mouse VIFs (see above).

The resulting VIF 3D structure reached a resolution of 7.2 Å (**Extended Data Fig. 5C, Extended Data Table 3**). The helical filament is composed of 5 protofibrils (**Fig. 2A, Supplementary Video 3**), which are connected laterally by individual contact sites between the protofibrils (**Extended Data Fig. 6, A and B**). Surprisingly, the protofibrils are also connected via a fiber in the VIF lumen (Fig. 2, B **and** C).

**Figure 2.**
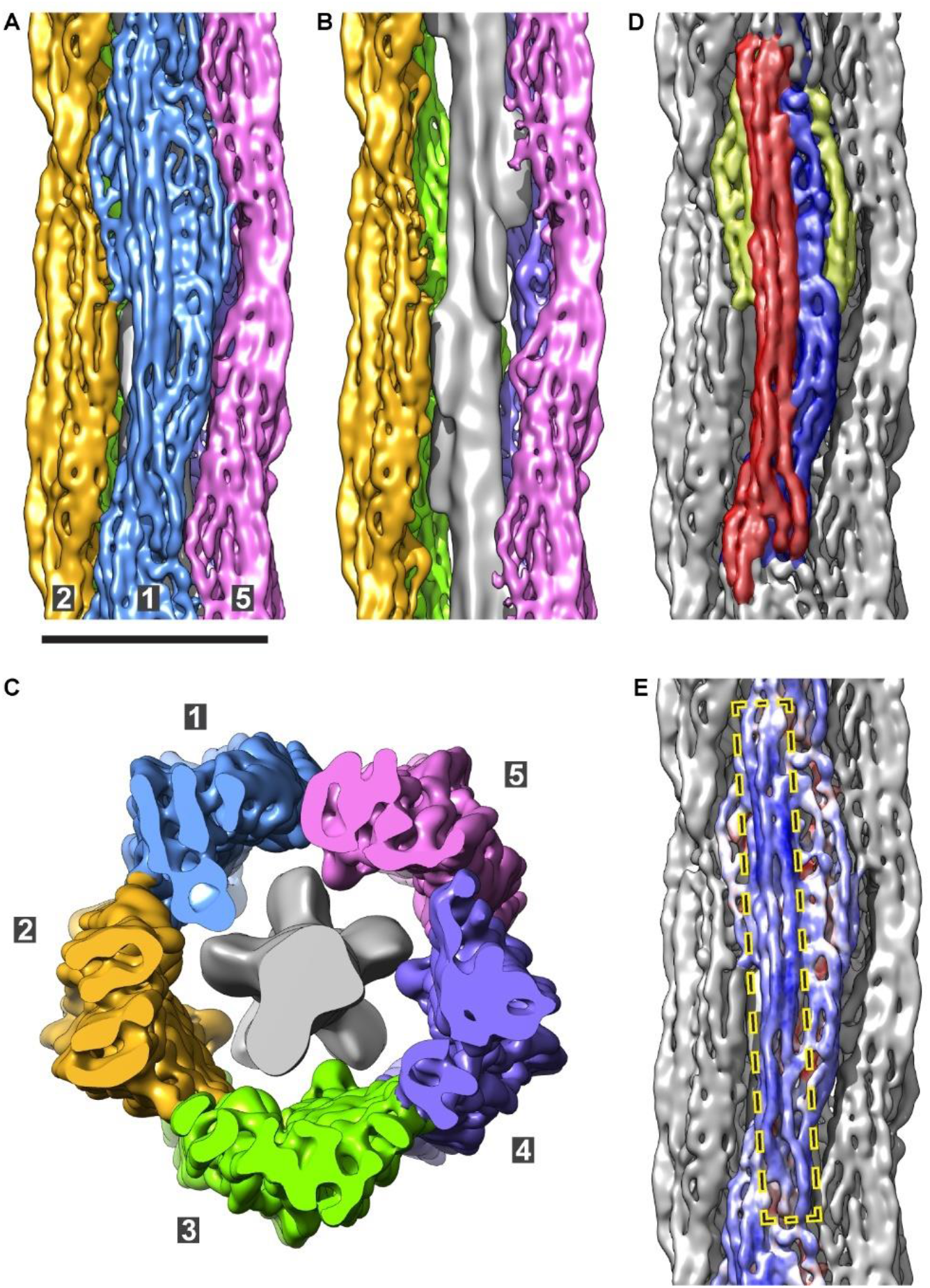
The 3D structure of VIFs. (**A**) Isosurface rendering of the VIF 3D structure. The protofibrils are shown in different colors and numbered from 1 to 5. Scale bar is 10 nm. (**B**) Omitting the front protofibril opens the view on the luminal fiber (grey density), which is located in the lumen of the VIFs. (**C**) Cross-section of VIFs displays all 5 protofibrils and the luminal fiber. (**D**) Segmentation of the repeating unit of a protofibril. The frontal, straight tetrameric α-helix bundle is colored in red, the lateral, curved tetrameric α-helix bundle in blue, and the contact sites between adjacent protofibrils in yellow. (**E**) The least flexible region of the repeating unit, indicated by mostly bluish colors, is the frontal, straight tetrameric α-helix bundle (dashed rectangle). Increased structural plasticity is indicated by reddish colors.

The repeating unit of a protofibril is assembled from three distinct regions (**Fig. 2D**). The first region is a relatively straight, tetrameric α-helix bundle (**Fig. 2D, red region**), that is ∼21 nm in length and slightly tilted with respect to the helical axis. This region forms large parts of the outer surface of the filament. Furthermore, it is the least flexible part of a protofibril (**Fig. 2E**). The second region is a more curved tetrameric α-helix bundle (**Fig. 2D, blue region**), which is attached laterally to the first bundle in its center and at its ends proceeds below the first bundle, eventually forming large parts of the inner surface of the filament, facing the lumen. Therefore, the central part of the repeating unit of a VIF protofibril is an octameric complex of 8 α-helices. Finally, the third region is split between both sides of the repeating unit and establishes lateral contact sites between adjacent protofibrils (**Fig. 2D, yellow regions**).

### Position of the tail domains

To localize the tail domains in the cryo-EM density map, we polymerized recombinantly expressed human vimentin lacking the tail domain [38], and applied the above cryo-EM workflow to the tailless VIFs. Surprisingly, the resulting VIF-ΔT structure clearly lacks the contact sites between the protofibrils (**Extended Data Fig. 6, C and D**), but the octameric repeating unit and the luminal fiber are of similar volume compared to the full-length VIF structure. Based on this result we conclude that the tail domains of vimentin, despite the fact that they are of low sequence complexity, are able to condense to a persistent structure in VIFs and form contacts between the protofibrils [57].

### Atomic model of VIFs

To build the atomic model of VIFs, based on our electron density map, we combined alphafold prediction [58] and molecular dynamics flexible fitting [59, 60], and used reported cross-linking mass spectrometry data as distance restraints [61-63]. As an additional restraint, we incorporated the position of the tail domains in the model building. The definition and initial folds of the vimentin domains, namely the 1A-1B (residues 86-253) and 2A-2B (265-412) domains, the linker L12 (254-264), and the head (1-85) and tail domains (413-466), were based on the alphafold prediction of the full-length vimentin dimer (**Extended Data Fig. 7**) [31]. These initial folds were adapted to the cryo-EM density map by molecular dynamics flexible fitting.

The resulting VIF 3D atomic model (**Fig. 3A, Extended Data Fig. 8, and Supplementary Video 4**) describes the cryo-EM density map in its entirety. The frontal, straight tetrameric α-helix bundle (**Fig. 2D, red region**) accommodates two antiparallel 2A-2B dimers (**Fig. 3B, Extended Data Fig. 8, green to red colored α-helices**). The C-terminal ends (CTEs) of each 2A-2B dimer position the tail domains to form the contact sites between the protofibrils (**Fig. 3B, Extended Data Fig. 8, magenta chains**), which are modelled containing a β-hairpin motif (**Extended Data Fig. 7**).

**Figure 3.**
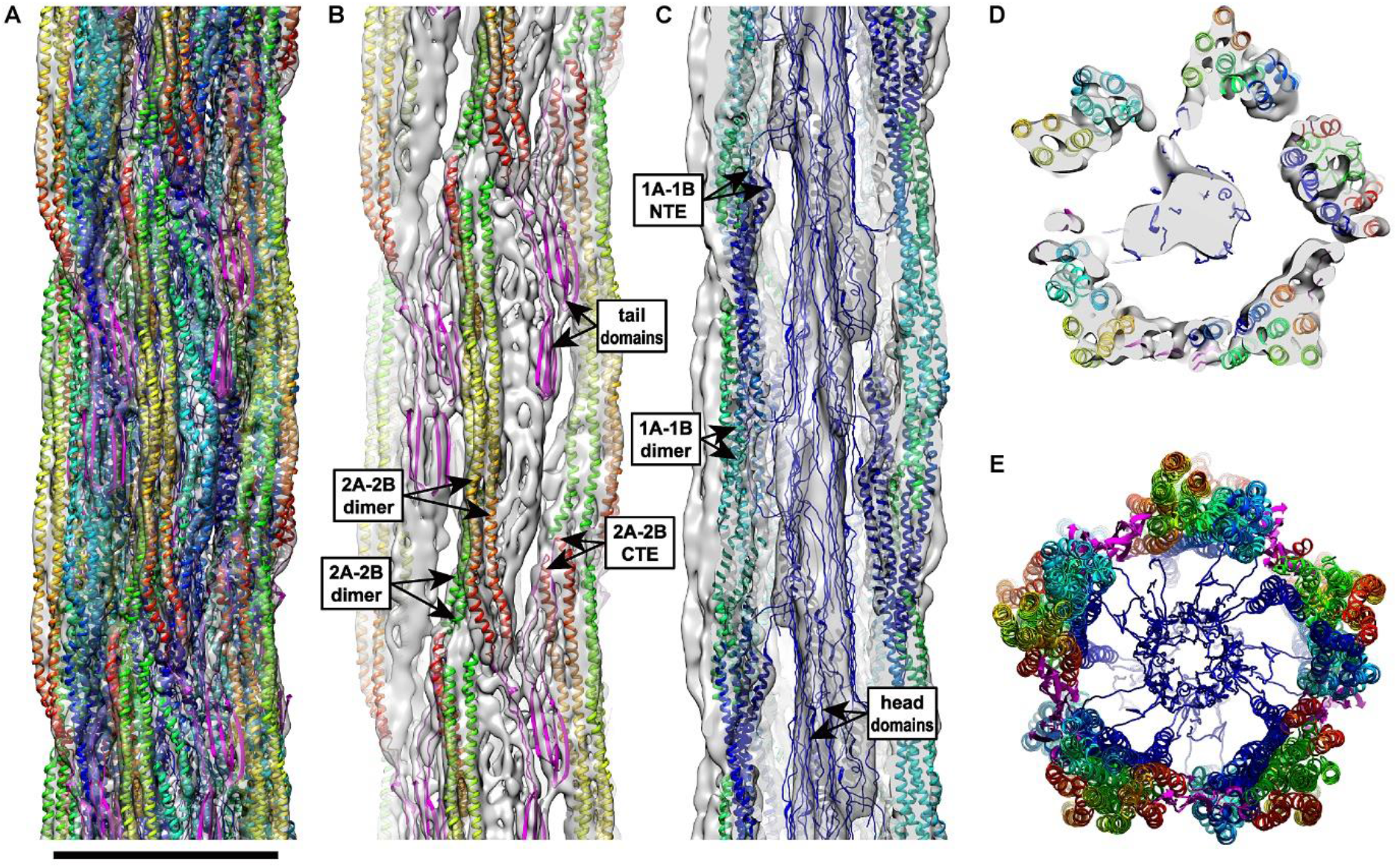
Atomic model of VIFs. (**A**) Complete VIF 3D atomic model docked into the density map (transparent grey). Scale bar is 10 nm. (**B**) The 2A-2B domains (α-helices colored from green**^NTE^** to red**^CTE^**) and the tail domains (magenta chains) are shown within the density map. Two antiparallel 2A-2B dimers form a straight α-helix bundle, which constitutes about one half of a protofibril. The CTEs of each 2A-2B dimer position the tail domains to form the contact sites between the protofibrils. (**C**) The 1A-1B domains (α-helices colored from blue**^NTE^** to mint green**^CTE^**) and the head domains (blue chains) are shown within the density map. Two antiparallel 1A-1B dimers compose a curved α-helix bundle, which approximately constitutes the second half of a protofibril. In the image the front protofibril is omitted to reveal the inside of the filament. The NTEs of each 1A-1B dimer protruding into the filament lumen, where the head domains aggregate to form the luminal fiber. (**D**) In VIFs there are 40 polypeptide chains in cross-section, assembled into 5 protofibrils. (**E**) The protofibrils interact laterally via the tail domains and centrally via the head domains. The 2A-2B dimers substantially shape the outer surface of VIFs, while the 1A-1B dimers predominantly coat their inner surface.

The lateral, curved tetrameric α-helix bundle (**Fig. 2D, blue region**) accommodates two antiparallel 1A-1B dimers (**Fig. 3C, Extended Data Fig. 8, blue to mint green colored α-helices**). The N-terminal ends (NTEs) of each 1A-1B dimer position the head domains to protrude into the filament lumen, where they interact to form the luminal fibre (**Fig. 3C, Extended Data Fig. 8, blue chains**). This finding is in agreement with previous work showing that the isolated, low complexity head domains of vimentin, and of other types of IFs such as neurofilaments, self-assemble into amyloid-like fibers, which are stabilized by characteristic cross-β interactions, as detected by X-ray diffraction [64].

In cross-section, VIFs comprise 40 polypeptide chains (**Fig. 3D, Supplementary Video 5**), which are partitioned into 5 octameric protofibrils. The protofibrils are connected laterally by interactions of the tail domains and connected centrally by interactions of the head domains (**Fig. 3E**). The 2A-2B dimers substantially shape the outer surface of VIFs, while the 1A-1B dimers predominantly coat the inner surface of the filaments.

### Modular assembly of VIFs

The vimentin tetramer (**Fig. 4A**) is the underlying asymmetric unit of the helical structure of VIFs. In general, the tetramer is straight extending ∼65 nm between CTEs of the flanking 2A-2B dimers. In particular, the α-helices of the 2A-2B dimers exhibit a mostly parallel geometry over a length of ∼21 nm while rotating ∼65° with respect to each other. In contrast, the four α-helices forming the central 1A-1B region exhibit a more pronounced coiled-coil geometry. This section is slightly curved, so that the NTEs of the 1A-1B dimers are protruding out of the plane of the tetramer and are positioned in pairs ∼29 nm apart. The head domains are proceeding parallel to the tetramer (within the luminal fiber in the mature filament). Our tetramer model predicts that the tail domains can exist in two conformations, either extending parallel to the long axis of the tetramer or folding back on one of the 2A-2B dimers. The possible range of motion of the tail domains is ∼10 nm.

**Figure 4.**
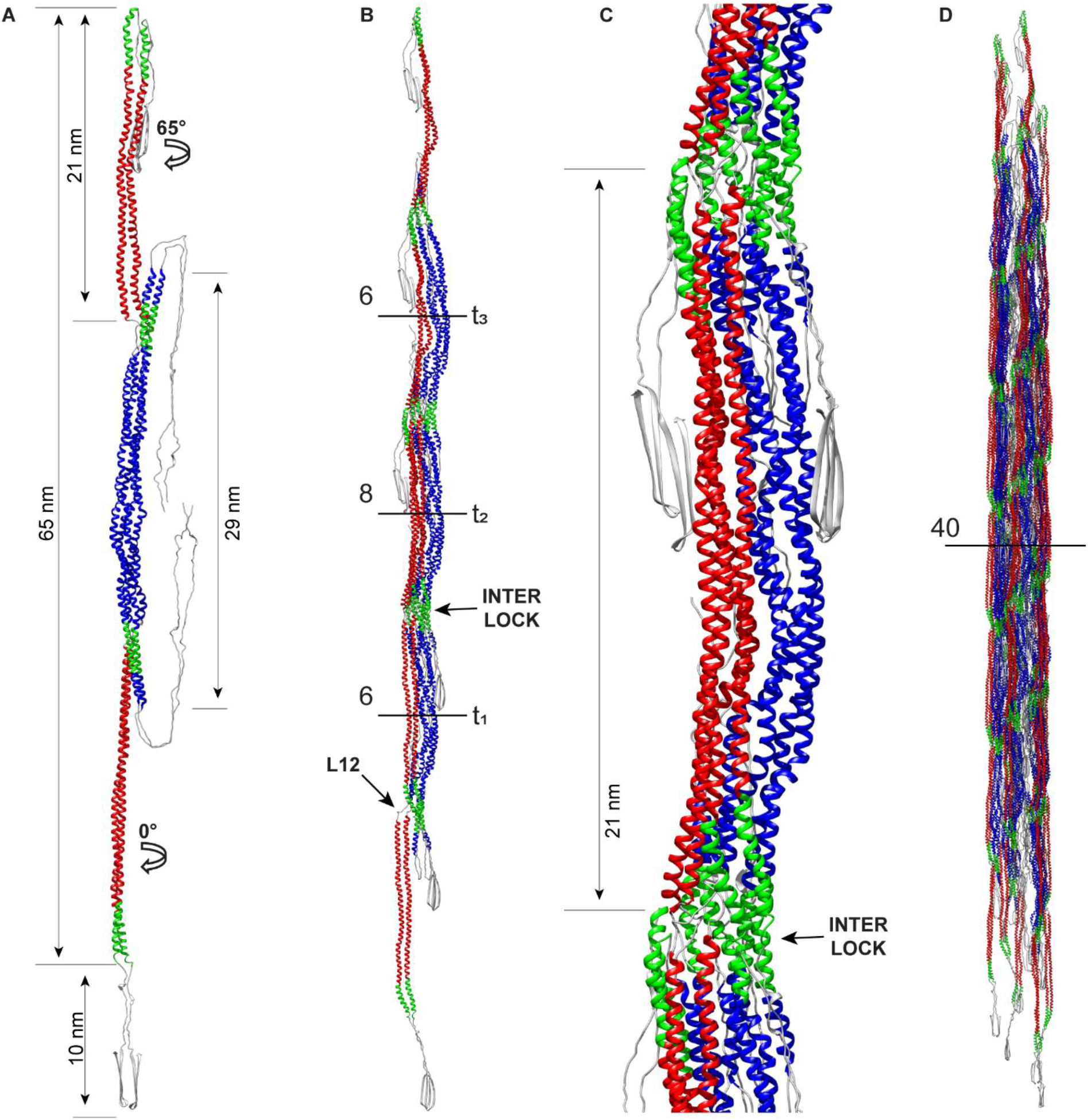
Building blocks of VIFs. (**A**) Tetramer model. Red colored α-helices are the 2A-2B dimers, blue colored α-helices are the 1A-1B dimers. Green colored α-helical segments are highly conserved regions between IFs in the 1A and 2B domains, respectively. (**B**) Protofibril model. The length is ∼110 nm between CTEs of the flanking 2A-2B dimers. Numbers and horizontal lines indicate the number of polypeptide chains in cross-section at the respective positions along the protofibril, and t_1_, t_2_, and t_3_ index the successive tetramers forming the visualized protofibril. (**C**) In the fully assembled repeating unit of a protofibril the interlock regions are formed on both sides of the assembly and are spaced ∼21 nm along the proto-fibrils. (**D**) Extended VIF model constructed from 30 tetramers with a length of ∼190 nm.

The full assembly of a protofibril with 8 polypeptide chains in cross section requires the sequential interaction of at least 3 tetramers (**Fig. 4B, Extended Data Fig. 9**). Starting from a first tetramer t_1_, a second tetramer t_2_ attaches with its 1A-1B section laterally to one of the flanking 2A-2B dimers of t_1_. This creates an intermediate assembly with 6 chains in cross section. Subsequently, a third tetramer t_3_ binds to either side of this assembly. This creates a minimal length (∼110 nm) protofibril with its basic repeating unit (8 chains in cross section) fully assembled in the center (**Fig. 4B**).

The construction of an interlocking mechanism in VIFs is intriguing. The 1A (residues 100-125) and 2B (residues 380-411) domains are highly conserved between IFs [29, 53]. Although they are distributed over the full length of the tetramer, in the fully formed protofibril they are brought into close proximity (**Fig. 4, green colored α-helical segments**), promoting interactions and interlocking successive tetramers together. This process is facilitated by the flexibility of the L12 linker domain, which allows the 2A-2B dimers to align parallel to the 1A-1B sections, and eventually find the right position to form the interlock region between the 1A and 2B domains. In the fully assembled repeating unit of a protofibril (**Fig. 4C**), both 2A-2B antiparallel dimers are laterally aligned with the 1A-1B section, and both interlock regions are fully formed ∼21 nm apart on either side of the assembly. The repeating unit unifies the three basic molecular interactions of vimentin dimers forming tetrameric units, defined as the A_11_, A_22_, and A_12_ binding modes, as shown by cross-linking mass spectrometry [61]. Mapping these cross-links on our model shows its excellent agreement with the original biochemical data measuring a mean distance between the cross-linked lysine residues of 9.4 Å (**Extended Data Fig. 10**) [63].

Finally, for a minimal length VIF model with 40 polypeptide chains per cross-section in its center the same rule applies, that at least 15 tetramers (3 per minimal length protofibril and 5 protofibrils) are needed to form such a fully polymerized VIF (**Fig. 4D, Extended Data Fig. 11, Supplementary Video 6**).

## Discussion

Advances in cryo-FIB/cryo-ET methodologies, in combination with extensive statistical analyses and single particle cryo-EM, have enabled us to visualize and describe the unprecedented molecular architecture of VIFs.

To obtain a physiologically relevant structure of VIFs, it was critical to measure their polymerization state in-situ. Here we show that VIFs are built from 5 protofibrils in cells. It was important to address this question, because of the significant structural polymorphism of VIFs [38, 40]. In this context, it was also important to consider that unprocessed tomograms have anisotropic resolution. Without correction, the difference between 4 and 5 protofibrils could not be resolved (**Extended Data Fig. 12**) [47].

We determined the helical symmetry of VIFs in-situ, which facilitated their 3D reconstruction. Based on a cryo-EM dataset of in-vitro polymerized VIFs we were then able to obtain a structure of fully assembled human VIFs. We have generated an atomic model of VIFs that maps the position and spatial relationships of the vimentin protein domains within the polymer and explains its unique building principles.

In comparison to F-actin and microtubules (**Fig. 5**), the basic building blocks of IFs are not globular proteins, but instead elongated, flexible tetramers. The tetramers intertwine during polymerization, complicating the structural characterisation of IFs, and partially concealing their helical architecture. Unlike F-actin and microtubules, the proteins forming IFs contain large stretches of low complexity sequence head and tail domains. Therefore, the structure of VIFs is substantially different, since VIFs integrate low complexity domains within a highly-ordered helical structure. This introduces an additional layer of structural complexity, based on transient molecular interactions [57, 64, 65].

**Figure 5.**
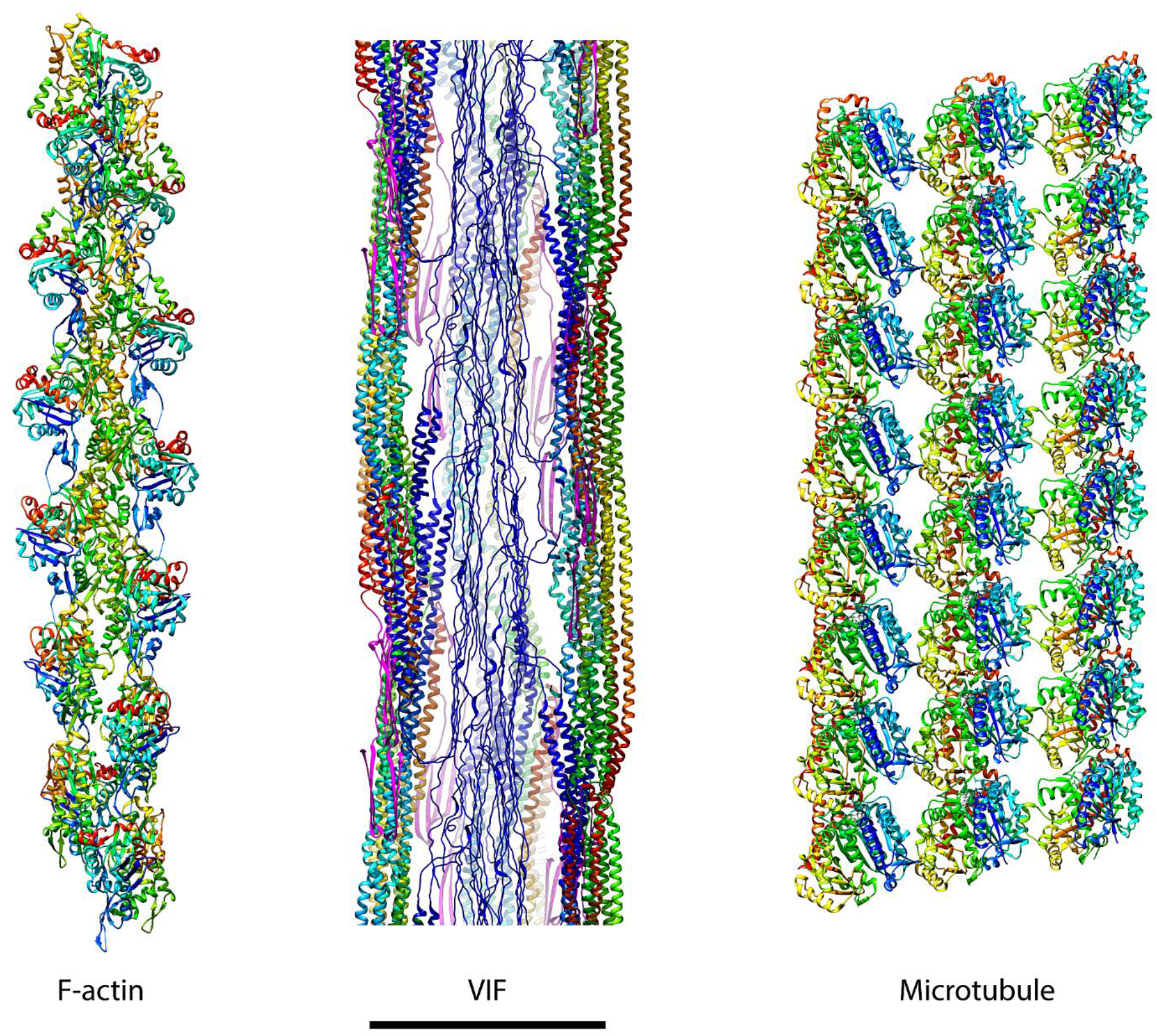
Complete structural picture of the cytoskeleton. The functions of the cytoskeleton in cells of mesenchymal origin are based on three biopolymers, which are depicted in this figure. F-actin and microtubules are built from compact, globular proteins, while VIFs are built from elongated, intertwining tetramers. All three filaments are assembled with helical symmetry. However, in VIFs a significant part of the structure is formed from low complexity sequence domains, which condense to connections between the protofibrils and form an amyloid-like fiber in the lumen of VIFs. This introduces an additional layer of structural complexity, based on transient molecular interactions. Three protofilaments of the microtubule are shown. Scale bar is 10 nm.

At the level of the individual protofibrils, classical molecular interactions (α-helices, coiled-coils) are prevalent. The lateral alignment of the 1A-1B and 2A-2B dimers and the formation of a molecular interlock region composed of the highly conserved 1A and 2B domains intertwine the tetramers to form a protofibril. However, at the level of the fully assembled filament, transient molecular interactions between intrinsically disordered protein domains play an important role. The protofibrils interact laterally via the tail domains while the head domains form an amyloid-like fiber in the lumen of VIFs.

We suggest that the combination of these molecular interactions results in the high elasticity and remarkable mechanical strength of VIFs [10-13]. Determining the relative contribution of these interactions to the mechanical properties of VIFs would be aided by the polymerization of headless VIFs. However, headless vimentin has been shown to not polymerize to VIFs [38]. This observation can be explained by our VIF model, since it is the head domains aggregating to the luminal fiber [64, 65]. This fiber may act as an anchor that centrally connects the protofibrils and keeps the filament together.

Considering the high sequence conservation in the 1A and 2B regions across IFs [29, 53], we hypothesize that the molecular interlocking mechanism is similar across IFs. The distance between the interlock regions is ∼21 nm. This axial repeat distance is also conserved across IFs [1, 30, 51-53]. It was shown that perturbing the interlocking mechanism with a single point mutation (Y117L) [66], prevents VIFs from elongating into to filaments. This observation can be explained by our VIF model, as without a functional interlocking mechanism the octameric repeating unit of VIFs cannot be formed.

The lateral alignment of the 1A-1B and 2A-2B dimers, the formation of a molecular interlock region, and the tail domain interactions between the protofibrils can only occur in VIFs at higher order polymerization states than the tetramer. However, the interactions of the head domains aggregating to the luminal fiber are possible at the tetramer stage. Therefore, shorter minimal length VIFs would also be stable (**Extended Data Fig. 11**). Such vimentin assemblies have been experimentally characterized and termed unit-length filaments [38, 39]. We suggest that a unit-length VIF is assembled from 5 tetramers and that this assembly is driven by a phase-separation process which facilitates the initial association of the tetramers via their head domains. Therefore, the head domains may act as initial nucleators of VIF filament assembly and elongation [64, 65].

This unit-length VIF would exhibit a length of ∼60 nm (**Extended Data Fig. 11**) [38]. Since the tail domain contact sites between the tetramers are not yet formed in this assembly, it would lack diameter control and tend to show larger diameters than that of VIFs. It has been shown experimentally that tailless VIFs lack diameter control [38] and that a compaction step occurs between unit-length filaments and VIFs [39]. We suggest that if the tail domain contact sites and octamer interlocks are formed, the VIF compacts to its final diameter of 11 nm.

Our results complete the structural picture of the cytoskeleton in cells of mesenchymal origin (**Fig. 5**). A comprehensive understanding of the mechanical properties of the cytoskeletal network cannot be achieved without first obtaining structural information for all of its components. With this structure of VIFs, it is now possible to take an integrative view of the cytoskeleton. Further understanding the unique utilization of low complexity domains in VIFs, as well as similar domains in other IFs, may begin to explain how IFs complement F-actin and microtubules to generate cell-type specific mechanical properties.

## Author contributions

M. E. conceived the research, analyzed all data, developed data analysis methods, built the VIF model, and wrote the manuscript. R. K-T. prepared the cryo-FIB samples and recorded the cryo-ET data. J. K. and S. K. provided the purified full-length vimentin, M. S. W. prepared the samples and recorded the cryo-EM data. R. B-P. cloned, produced, and polymerized the delta-tail vimentin and prepared the cryo-EM samples. C. T. B. recorded the cryo-EM data of the delta-tail vimentin and edited the manuscript. Y. T. prepared the detergent-treated MEF samples and recorded the cryo-ET data. S. S. prepared MEF samples and recorded the light microscopy data. R. D. G. conceived the research and edited the manuscript. O. M. conceived the research, supervised the project, edited the manuscript and acquired the funding.

## Data availability

The VIF structure is deposited in the Electron Microscopy Data Bank under the accession code EMD-16844. The VIF atomic model will be deposited into the PDBDEV upon publication.

## Acknowledgements

This work was funded by grants from the Swiss National Science Foundation (SNSF 310030_207453) and the Mäxi Foundation to O.M. We thank the Center for Microscopy and Image Analysis at the University of Zurich. R. D. G. and S. S. were supported by an NIH program project grant awarded to R. D. G.

## Methods

### Cell lines and cell culture

MEFs were cultured in DMEM (Sigma-Aldrich, D5671), supplemented with 10% FCS (Sigma-Aldrich, F7524), 2 mM L-Glutamine (Sigma-Aldrich, G7513) and 100 μg/ml penicillin/ streptomycin (Sigma-Aldrich, P0781), at 37°C and 5% CO_2_ in a humidified incubator.

### 3D-SIM imaging

Sub-confluent cultures of MEFs growing on #1.5 glass coverslips were fixed with 4% paraformaldehyde for 10 min at RT. The fixed cells were permeabilized with 0.1% Triton-X 100 for 10 min at RT and then incubated with chicken anti-vimentin (1:200, 919101, Biolegend, CA, USA) for 30 min in phosphate buffered saline (PBS) containing 5% normal goat serum. This was followed by staining with goat anti-chicken Alexa Fluor 488 (1:400, A-11039, Invitrogen, CA) and DAPI in PBS for 30 min. The stained cells were mounted with ProLong Glass Antifade Mountant (Life technologies, Carlsbad, CA, USA).

Moreover, MEFs were seeded on #1.5 glass coverslips and the next day they were washed with PBS/2 mM MgCl_2_ for 5 s followed by incubation with PBS containing 0.1% Triton X-100, 10 mM MgCl_2_, 0.6 M KCl and protease inhibitors for 25 s at RT. The extracted cells were rinsed with PBS/2 mM MgCl_2_ for 10 s and subsequently incubated with 2.5 units/µl Benzonase for 30 min at RT. After rinsing with PBS/2 mM MgCl_2_ the cells were fixed with 4% paraformaldehyde for 5 min at RT. The fixed cells were then stained with chicken anti-vimentin (1:200, 919101, Biolegend, USA) and rabbit anti-lamin A (to determine the location of the nucleus) in PBS containing 5% normal goat serum for 30 min at RT followed by incubation with goat anti-chicken and anti-rabbit secondary antibodies (1:400, A-11039, A-11011, Invitrogen, USA) for 30 min at RT. After washing in PBS, the stained cells were mounted with Prolong Glass Antifade Mountant (Life technologies, Carlsbad, CA, USA).

3D-SIM imaging was carried out with an N-SIM Structured Illumination Super-resolution microscope system (Nikon, Tokyo, Japan) using an oil immersion objective lens (SR Apo TIRF100X, 1.49 NA, Nikon). For 3D SIM, 26 optical sections were imaged at 100 nm interval. The raw SIM images were reconstructed with the N-SIM module of Nikon Elements Advanced Research with the following parameters: illumination contrast, 1.00, high-resolution noise suppression, 0.75, and out-of-focus blur suppression, 0.25. Brightness and contrast were adjusted for image presentation.

### Cryo-FIB/Cryo-ET in-situ analysis of VIFs

MEFs were seeded onto glow-discharged, carbon-coated, gold EM grids (Quantifoil, Jena, Germany, holey carbon R 2/1, Au 200) in standard medium overnight. After a single wash in 1X PBS (Fisher bioreagents, BP399-1) the grids were vitrified in liquid ethane using a manual plunge freezing device.

The grids were coated with 5-10 nm Pt/C with a Leica BAF060 system cooled to -160°C, prior to cryo-FIB milling. Next, the grids were transferred to the FIB-SEM (Zeiss Auriga 40 Crossbeam), equipped with a Leica cryo-stage. After transfer, the grids were coated with two 5 s flashes of organometallic platinum using the internal gas injection system set to 28°C. The cells were milled with a focused gallium ion beam at a constant voltage of 30 kV and a current between 10 and 240 pA at a stage angle of 18°. The process was controlled by the Nano Patterning and Visualization Engine (NPVE, Zeiss) and monitored with the SEM at 5 kV. The resulting FIB milled lamellae were 100-200 nm thick (**Extended Data Fig. 1A**).

The cryo-FIB milled grids were transferred to a Titan Krios 300 kV cryo-TEM (Thermo Fisher Scientific, Waltham, USA) equipped with a K2 summit direct electron detector and a Quantum energy filter (Gatan, Pleasanton, USA). The tilt series were acquired with SerialEM [67] using a 100 µm objective aperture at a magnification of 64,000x, at a pixel size of 2.21 Å, with -4 µm defocus and using a dose-symmetric tilt scheme [68] from -60° to +60° in 3° increments starting from 0°. Overall, 102 tilt series were acquired, each with a total dose of ∼160 e^-^/A^2^.

Deconvolution of missing wedge induced distortions in the cryo-FIB tomograms was performed with IsoNet [46]. A subset of seven tomograms was selected from the cryo-FIB dataset and further processed with IsoNet. The selected tomograms contained a tilted lamella showing a region around the nuclear membrane, with lamin filaments, chromatin, NPCs, actin filaments, MTs, VIFs (**Fig. 1, B and C**), and platinum depositions in varying sizes on the surface of the lamellas (**Extended Data Fig. 1A, yellow dashed rectangle**).

Initially, the tomograms were boxed to center the lamella and 4x binned to a size of 960 x 928 x 360 voxels, with a voxel size of 8.84 Å. For CTF correction, a signal-to-noise ratio fall off parameter of 0.7 was used. For automatic mask creation a density percentage of 50% and standard deviation percentage of 50% was chosen. Training of the missing wedge deconvolution neural network was based on randomly extracted 2000 sub-tomograms (box size 64^3^ voxels) per tomogram within the automatically created mask, with the following training parameter: 30 iterations, learning rate 0.0004, and drop-out 0.5. The displayed cross-sections of VIFs (**Fig. 1D, Extended Data Fig. 1B**) were extracted from one missing wedge corrected tomogram.

In order to model the influence of the missing wedge induced resolution anisotropy on the representation of the protofibrils in the tomographic cross-sections, we convolved the VIF structure with a missing wedge, reflecting a tilt range from -60° to +60°, and compared its cross-sections with cross-sections from the unaffected structure (**Extended Data Fig. 12**). The model calculation was performed using MATLAB scripts derived from the TOM toolbox [69].

### Cryo-ET of detergent-treated MEFs

MEFs were grown to ∼80% confluency on glow-discharged holey carbon EM grids (R2/1, Au 200 mesh; Quantifoil, Jena, Germany) prior to preparation for cryo-ET analysis. Grids that showed a relatively homogenous distribution of cells were selected using fine tweezers, and then washed in PBS/2 mM MgCl_2_ for 5 s. The grids were treated for 20-40 s in pre-permeabilization buffer (PBS containing 0.1% Triton X-100, 10 mM MgCl_2_, 600 mM KCl and protease inhibitors) and then rinsed in PBS/2 mM MgCl_2_ for 10 s. Next, the grids were incubated with Benzonase (2.5 units/µl in PBS/2 mM MgCl_2_; Millipore, Benzonase Nuclease HC, Purity >99%) for 30 min at RT. After washing the grids with PBS/2 mM MgCl_2_, a 3 µl drop of 10 nm fiducial gold markers (Aurion) was applied to the grids. For vitrification the grids were manually blotted for ∼3 s from the reverse side and plunge frozen in liquid ethane.

Tilt series acquisition was conducted using a Titan Krios transmission electron microscope equipped with a K2 Summit direct electron detector and Quantum energy filter. The microscope was operated at 300 keV with a 100 µm objective aperture. In total 225 tilt series were collected at a nominal magnification of 42,000x and the slit width of the energy filter was set to 20 eV. Super-resolution movies were recorded within a tilt range from -60° to +60° with 2° increments using SerialEM [67]. The image stacks were acquired at a frame rate of 5 fps with an electron flux of ∼2.5 e^-^/pixel/s. The tilt series were recorded with a total electron dosage of ∼125 e^-^/Å^2^ and within a nominal defocus range between -2 μm to -6 μm. The super-resolution image stacks were drift-corrected and 2x binned using MotionCorr [70], resulting in a pixel size of 3.44 Å for the tilt series. For each projection the defocus was measured and the contrast transfer function was corrected by phase-flipping. Then, from each tilt series a 4x binned overview tomogram was reconstructed (**Extended Data Fig. 2B**). CTF correction and tomogram reconstruction was performed using MATLAB scripts (MathWorks, Natick, USA) derived from the TOM toolbox [69, 71].

Algorithms from EMAN2 [72] were employed for training a convolutional neural network capable of segmenting VIFs in the overview tomograms. The segmentations were manually checked and cleaned from obvious false positive VIF detections in Chimera [73]. Based on scripts derived from the ActinPolarityToolbox (APT) [48], two sets of segment coordinates were extracted from the segmentations. In the first set (second set, respectively) the picking distance along the VIFs was set to 165 Å (55 Å), resulting in 390,297 (1,148,072) segment coordinates. Next, based on the segment coordinates two stacks of subtomograms were reconstructed from the CTF corrected tilt series with the TOM toolbox. The dimensions of the subtomograms were 65 x 65 x 65 nm^3^ and 38 x 38 x 38 nm^3^, respectively, and the voxel size was 3.44 Å. Subsequently, APT scripts were applied to project the subtomograms, using a projection thickness of 331 Å for the first set and 220 Å for the second set. The size of the subtomogram projections derived from the first and second coordinate sets were 65 x 65 nm^2^ and 38 x 38 nm^2^, respectively.

### Initial estimate of helical symmetry

The VIF segments were subjected to multiple rounds of unsupervised 2D classifications in RELION [49, 55]. 2D classes which were not combining single VIFs (for example, segments containing multiple VIFs running parallel or crossing on top of each other) or containing false-positive VIF detections (for example, actin filaments or vesicle membranes) were excluded based on visual inspection. As a consequence, the first particle set (second particle set, respectively) was concentrated to 133,780 (615,106) VIF segments.

Based on the resulting 2D class averages from the first set of particles (**Fig. 1E**), 2D auto-correlation functions of the class averages were calculated with the TOM toolbox function *tom_corr*, then displayed as profile plots and averaged (**Extended Data Fig. 2C**), in order to measure the distance between similar features observed in the class averages (**Fig. 1E, yellow asterisks**). The resulting 2D class averages from the first particle set were also used to calculate an averaged power spectrum with the TOM toolbox function *tom_ps* in order to prove the helical symmetry of VIFs and to obtain an estimate of their helical symmetry parameter (**Fig. 1F**). This initial estimate was 72° for the helical twist angle and 37 Å for the helical rise.

### Computational filament assembly

The assembly of extended stretches of VIFs (computationally assembled VIFs, ca-VIFs) was based on the 2D classification of the second particle set (**Extended Data Fig. 3A**). For this purpose, the 2D transformation calculated for each segment (namely its in-plane rotation angle and xy-translation) was inverted and applied to the respective class averages, so that the inversely transformed class averages match position and orientation of the segments in the tomogram image frame [4, 74]. As a result of this operation the ca-VIFs are represented by a series of class averages (**Extended Data Fig. 3, B and C**), which significantly improves their signal-to-noise ratio compared to the raw filaments. Additionally, the ca-VIFs were unbent (**Extended Data Fig. 3D**) based on a MATLAB algorithm derived from the ImageJ [75] straighten function [1, 76].

In total, 5205 ca-VIFs of different lengths were assembled (**Extended Data Fig. 4A**). We selected a subgroup of 389 ca-VIFs with a length of ≥353 nm. These were boxed to equal length (353 nm) and then measured employing autocorrelation (**Extended Data Fig. 4B**), whether the previously detected periodicity (**Fig. 1E, yellow asterisks**) persists over much longer distances. This result showed a long-range periodic pattern in 2D projections of VIFs that repeats every 186.5 Å ± 26.0 Å, supporting the previous measurements on single class averages.

### Determination of helical symmetry

Next, we calculated a combined power spectrum of the ca-VIFs (**Extended Data Fig. 4C**). Due to the increased resolution, the previously detected layer lines (**Fig. 1F**) are splitting in fine layer lines and a dense spectrum of layer lines is revealed. For determination of the helical symmetry of VIFs based on the power spectrum of the computationally assembled filaments, the following procedure was developed in MATLAB.

Firstly, 1-pixel wide rows in the interval between 1/30 Å and 1/69 Å were sequentially extracted from the power spectrum and compared by cross-correlation with a zero order Bessel function (assuming a filament radius of 55 Å). By this means the similarity of the extracted rows with a meridional reflection was measured, which indicates the helical rise of the underlying helical assembly [50]. As a result, the layer line at 1/42.5 Å was identified as related to the helical rise of VIFs (**Extended Data Fig. 4D**).

In the next step, the sequence of layer lines (**1/207.4 Å, 1/195.9 Å, 1/185.6 Å, 1/176.3 Å; shown in Extended Data Fig. 4C, inset**), which are organized around the layer line that reflects the long-range periodic pattern found previously (**Extended Data Fig. 4B**), were related to a similar sequence of layer lines found around the meridional reflection (**1/42.5 Å, 1/40.1 Å, 1/37.9 Å, 1/36.3 Å; shown in Extended Data Fig. 4D**). To this end, the relationship n=P/h_r_ was applied that connects the number of asymmetric units (n) with the helical pitch (P) and helical rise (h_r_) of a helical assembly. In particular, the optimal n was searched that relates the two given sequences of layer lines by P=n•h_r_, interpreting the first sequence as layer lines associated with the helical pitch of VIFs and the second sequence as layer lines associated with the helical rise of VIFs. The result of this calculation was n=4.8824 (**Extended Data Fig. 4E**). In summary, a helical twist angle of 73.7° (that is 360°/n) and a helical rise of 42.5 Å were determined based on the power spectrum of ca-VIFs.

However, due to the high density and complexity of the layer line spectrum we were not able to index the Bessel peaks [50], so that the uniqueness of this solution must be proven by other means. Therefore, we conducted extensive helical 3D classifications and helical symmetry searches based on single particle data, which was converging independently to a numerically identical solution (see next section).

### Cryo-EM of in-vitro polymerized human VIFs

Human full-length vimentin was expressed in a transformed BL21 *E. coli* strain and isolated from inclusion bodies as described previously [77]. The protein was stored in 8 M urea, 5 mM Tris HCl (pH 7.5), 1 mM EDTA, 10 mM methylamine hydrochloride (Merck, Munich, Germany) and approximately 0.3-0.5 mM KCl at -80°C.

To reassemble human vimentin, it was subjected to a stepwise dialysis in phosphate buffer at pH 7.5. Therefore, a dialysis tube with a 12-14 kDa cutoff (Serva) was rinsed three times with dialysis buffer 1 (6 M urea, 2 mM phosphate buffer). Next, 200 µl of purified vimentin at 0.2 mg/ml in 8 M urea was added to the dialysis tube and dialyzed for 1 h at room temperature against dialysis buffer 1 under gentle magnetic mixing. The buffer was exchanged three times in 1 h intervals against fresh dialysis buffer with decreasing concentrations of urea (dialysis buffer 2: 4 M urea, 2 mM phosphate buffer, dialysis buffer 3: 2 M urea, 2 mM phosphate buffer, dialysis buffer 4: 1 M urea, 2 mM phosphate buffer). In the last step, the buffer was exchanged against 2 mM phosphate buffer and vimentin was incubated for 2 h at room temperature.

VIF formation was initiated by dialyzing vimentin against 2 mM phosphate buffer containing 100 mM KCl at pH 7.5 at 37°C overnight. To prepare samples for cryo-EM, 4 µl of filamentous vimentin at a concentration of 0.1 mg/ml was applied to glow-discharged, carbon coated copper grids (Cu R2/1, 200 mesh, Quantifoil) and vitrified in liquid ethane using a manual plunge freezing device.

The VIF grids were imaged using a 300 kV Titan Krios G3i cryo-electron microscope (Thermo Fisher) equipped with a K3 direct electron detection camera (Gatan) mounted on a Bio Quantum Energy Filter (Gatan). Dose-fractionated micrographs were acquired with a 100 µm objective aperture and zero-loss energy filtering using a 20 eV slit width, at a magnification of 130,000x with a pixel size of 0.34 Å in super-resolution mode using EPU (Thermo Fisher). A defocus range between -0.8 and -2.8 μm was chosen. The frame exposure time was set to 0.013 s with a total exposure time of 1 s per (75 frames in total), resulting in a total electron dosage per dose-fractioned micrograph of ∼62 e^-^/Å^2^. The micrographs were corrected for beam-induced motion with RELION [55] and binned by Fourier cropping, resulting in 12,160 micrographs with a pixel size of 0.68 Å. For CTF estimation Gctf [78] was used. In particular, the estimated maximal resolution per micrograph histogram was optimized as a function of the amplitude contrast fraction parameter. This procedure yielded an improved CTF model to describe the data with an amplitude contrast fraction of 0.2, as compared to the default value of 0.1.

### Single particle reconstruction of VIFs

Filament picking was performed with crYOLO [79]. To this end we used a neural network trained on in-vitro polymerized keratin K5/K14 IFs [4], which were acquired with identical cryo-EM microscopy parameter as the VIFs, and applied this neural network to the VIF micrographs. In the prediction step the filament mode was activated, with an initial box distance between the VIF segments of 27.2 Å. As directional method convolution was used, specifying a search range factor of 1.41 and a filament and mask width of 14 nm. As a result, 130,094 VIFs were picked from the micrographs, which were subdivided in 1,462,717 VIF segments with a particle box size of 38 x 38 nm^2^.

For initial particle sorting and cleaning the segments were extracted with a pixel size of 6.8 Å and subjected to multiple rounds of unsupervised 2D classifications in RELION [55, 56]. In between the 2D classifications, those 2D classes combining segments of low quality (for example, segments containing overlapping or unravelling VIFs or carbon edges) were excluded based on visual inspection, therefore reducing the number of particles gradually to 801,585 good VIF segments. At this stage the 2D classes (**Extended Data Fig. 5A**) already showed the above-described characteristic pattern that one filament wall appears more pronounced in projection than its counterpart (**Fig. 1E, Extended Data Fig. 2D, and Extended Data Fig. 3A**). However, at the same time the 2D classes showed that VIFs exhibit a pronounced variability regarding their diameter, similar to what we previously observed for keratin K5/K14 IFs [4].

Since the diameter of a helical filament is coupled to its helical symmetry, multiple rounds of helical 3D classifications were performed in RELION [55, 56] with the aim to reduce the heterogeneity in the particle set. As initial 3D template for helical 3D classifications a rotational symmetric filament tube was reconstructed from the VIF segments using 90° as approximation for the tilt angle, the refined psi angle from the previous 2D classification step, and a random angle as the rot angle [4]. As starting point for the helical symmetry search an initial twist angle of 72° and an initial helical rise of 37 Å was set (**Fig. 1F**), and this helical symmetry was also imposed on the initial 3D template. The actual helical symmetry search was performed in an interval between 50° to 100° for the helical twist and 30 Å to 57 Å for the helical rise, therefore also capturing possible helical assemblies based on 4 or 6 protofibrils.

In between the helical 3D classifications, those 3D classes converging to the borders of the search interval were removed, therefore reducing the number of particles gradually to 520,902 good VIF segments. In the final helical 3D classification, the mean helical symmetry calculated from all the VIF 3D classes was 73.4° for the helical twist angle and 42.1 Å for the helical rise. This helical symmetry parameter were used as the initial twist and rise values for subsequent local helical symmetry searches during 3D refinement.

In the following the processing of the particles was iteratively switched between 3D auto-refine and 3D classification jobs in RELION [55, 56]. VIF segments at this stage were extracted with a pixel size of 1.0 Å. For 3D auto-refine jobs the angular sampling was reduced to 1.8°, with local angular searches starting from 0.9°. Local helical symmetry search was switched on. Subsequently, a 3D classification job without image alignment was conducted. The 3D classes which were carried over to the next 3D auto-refine job were selected based on visual inspection. Additionally, particles converging to a distance <12 Å between neighbouring segments were removed. Thereby, the number of particles was gradually reduced to a final number of 236,920 good VIF segments. The final local helical symmetry search converged to a helical twist angle of 73.7° and a helical rise of 42.5 Å, which is numerically identical to the helical symmetry previously determined by power spectrum analysis (**Extended Data Fig. 4**). In order to improve the resolution of the VIF structure a reference mask was applied during the subsequent 3D auto-refine runs. This mask was based on an intermediate VIF structure, that was lowpass filtered to 30 Å. In this structure the luminal fibre was removed using UCSF Chimera [73], with the aim to reduce the structural heterogeneity in the averaging volume. The RELION command *relion_mask_create* was used to extend the binary reference mask by 6 voxels and add a soft edge of 12 voxels. The result of this masked 3D auto-refine job with applied helical symmetry (helical twist 73.7°, helical rise 42.5 Å) was sharpened with LocalDeblur [80]. The LocalDeblur parameter ʎ and K were set to 1.0 and 0.025, respectively, and the required local resolution map was calculated with ResMap [81].

For the final 3D auto-refine job the above reference mask was kept constant and the above sharpened structure was used as initial reference. However, in contrast to previous 3D auto-refine jobs, helical symmetry was relaxed and not applied during final 3D refinement. The resulting VIF 3D structure reached a resolution of 7.2 Å (**Extended Data Fig. 5C**).

To visualize the increased homogeneity of the VIF segments that were used in the final 3D auto-refine job, they were combined with a 2D classification job in RELION. Here the 2D alignment of the particles was fixed to the translations and in-plane rotations that were calculated in the final 3D auto-refine job (**Extended Data Fig. 5B**).

The final VIF 3D structure (**Fig. 2**) was sharpened with LocalDeblur (ʎ=1.0, and K=0.025), helical symmetry was imposed (helical twist 73.7°, helical rise 42.5 Å), and the structure was lowpass filtered to 7.2 Å. The local resolution distribution of the map was calculated with ResMap (mean resolution of all analysed voxels was 5.2 Å) and interpreted as a measure for local structural plasticity (**Fig. 2E**).

### Cryo-EM of in-vitro polymerized human VIFs-ΔT

The gene encoding human vimentin ΔT (amino acids 1-410) was cloned in pEt24d(+) and the protein was expressed in Rosetta2 pLysS as inclusion bodies [77]. The protein was purified in multiple steps consisting of isolation of the inclusion bodies, guanidinium chloride-based solubilisation, and size-exclusion chromatography of the recombinant ΔT vimentin. Briefly, the cell extract obtained after sonication of the frozen bacteria pellet in a buffer containing Tris HCl 50 mM pH 8, NaCl 200 mM, Glycerol 25%, EDTA 1 mM, lysozyme 10 mg/ml, MgCl2 20 mM, DNase 1 8 ug/mL, RNase A 40 ug/mL, NP40 1%, Deoxycholic Acid sodium 1%, one cOmplete™ Protease Inhibitor Cocktail (Roche), was centrifuged for 30min at 12,000g, at 4^◦^C. The inclusion bodies pellet was washed 3 times (or more if persistence of dark material) using cycles of resuspension/centrifugation (30min at 12,000g, at 4^◦^C) and a buffer containing Tris HCl 10 mM, pH 8, Triton X-100 0.5 %, EDTA 5 mM, DTT 1.5 mM, one cOmplete™ Protease Inhibitor Cocktail tablet. As a last step the inclusion bodies pellet was washed in a buffer containing Tris HCl, pH 8 10 mM, EDTA 1 mM, DTT 1.5 mM, one cOmplete™ Protease Inhibitor Cocktail tablet. The vimentin ΔT protein was then solubilised with GuHCl 6M, in 10 mM TrisHCl pH 7.5, and clarified for 30 min at 10,000g, at 4^◦^C. The vimentin-containing supernatant was collected. A size-exclusion chromatography using Superdex 200 Increase 10/300 GL (Cytiva) was performed. Protein purity was checked using SDS-PAGE.

The protein concentration was adjusted to 0.2-0.4 mg/ml, and VIF-ΔT filaments were reconstituted by serial dialysis of 30 min at 22°C using buffers of decreasing concentration of GuHCl, followed by an overnight dialysis step at 4°C in GuHCl free buffer. The dialysis buffers were 5 mM Tris HCl pH 7.5, with 1 mM EDTA, 0.1 mM EGTA, 1 mM DTT, containing 4, 2, 0 M GuHCl. The dialysed protein solution was used for VIF-ΔT assembly by dialysis using a high salt-buffer containing 10 mM Tris HCl pH 7.5 with 100 mM KCl.

To prepare samples for cryo-EM, 3 µl of the VIF-ΔT solution was applied onto glow-discharged holey carbon EM grids (Cu R2/1, 200 mesh, Quantifoil), which were subsequently blotted manually and plunge frozen in liquid ethane.

For imaging the VIF-ΔT grids the same EM setup was used as before, with the only difference that the dose-fractionated micrographs were recorded in counting mode. The micrographs were corrected for beam-induced motion with MotionCor2 [82] without binning, resulting in 19,534 micrographs with a pixel size of 0.68 Å. For CTF estimation Gctf [78] was used, with the same amplitude contrast fraction as for the CTF estimation of the VIF dataset.

### Single particle reconstruction of VIFs-ΔT

Filament picking was performed with crYOLO [79], with the same neural network and parameter settings as for the VIF dataset. In the VIF-ΔT dataset 1,019,393 VIF segments were initially picked with a particle box size of 38 x 38 nm^2^. Particle sorting and cleaning of the segments was performed as before with the VIF dataset, based on multiple rounds of unsupervised 2D classifications in RELION [55, 56] and low-quality particle exclusion based on visual inspection of 2D classes, therefore reducing the number of particles gradually to 143,270 good VIF-ΔT segments.

The VIF-ΔT structure was calculated in RELION with a 3D auto-refine job, without applying helical symmetry. The resulting structure reached a resolution of ∼14 Å. For visualization helical symmetry was imposed (helical twist 73.7°, helical rise 42.5 Å), and the structure was lowpass filtered to 14 Å (**Extended Data Fig. 6D**).

#### Model building

In the first stage, alphafold [58] was used to predict the model of a tetramer consisting of two 1A-1B chains and two 2A-2B chains. Two copies of this tetramer were docked by using ClusPro [83] to form an initial octameric model of the repeating unit of a VIF protofibril. The previously published cross-links [61, 62] were mapped [63] on this model and the distance between the cross-linked lysine residues were minimized by a simple geometric rotation and translation of the dimeric entities (namely the 1A-1B and 2A-2B dimers).

In detail, one 2A-2B dimer was rotated -5° around its center and translated +20 Å in z-direction, the second was rotated +5° around its centre and translated -20 Å in z-direction. Then, one 1A-1B dimer was translated -20 Å in z-direction and the second +20 Å in z-direction. These simple geometric transformations of the 1A-1B and 2A-2B dimers relative to each other improved the mean distance between the cross-linked residues to 15.6 Å.

Next, this cross-link corrected model was rigid body docked to the VIF 3D structure using UCSF Chimera [73] and then adapted to the electron density map by molecular dynamics flexible fitting using namdinator [60, 84].

Subsequently, the helical symmetry of VIFs was applied to form a model of a protofibril. From this model an initial tetramer model was extracted and converted to a simulated electron density map, which was used in combination with molecular dynamics flexible fitting to transform two copies of the alphafold model of the full-length vimentin dimer (**Extended Data Fig. 7**) to the overall shape of the tetramer. This initial complete tetramer model (including the head, tail, and linker L12 domains) was then further refined with the VIF electron density map using molecular dynamics flexible fitting, resulting in the final tetramer model (**Fig. 4A**). To assemble the complete VIF atomic model (**Fig. 3**) the helical symmetry of VIFs was applied to the final tetramer model.

### Visualization

All isosurface representations of the VIF structures and visualizations of the VIF atomic model were rendered with UCSF Chimera [73]. Protofibril segmentation and the segmentation of the repeating unit of a protofibril was conducted with Segger [85]. For the visualization of the complete cytoskeleton, PDB-8A2R [86] was used for the actin filament, and PDB-6DPU [87] for the MT.

## Extended Data Figures

**Extended Data Figure 1.**
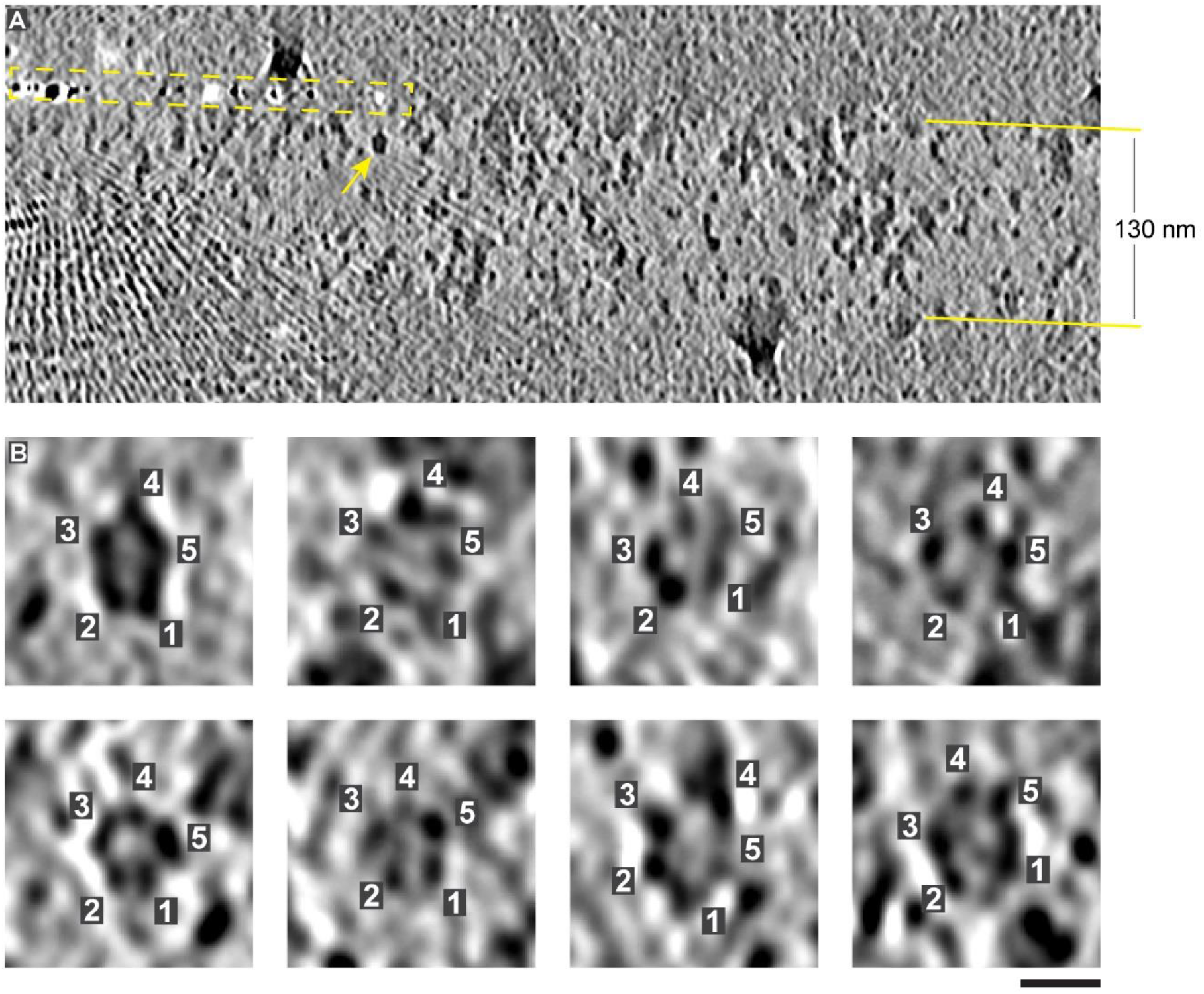
Cross-section analysis of VIFs. (**A**) Slice in x-z-direction with 8.84 Å thickness through a tomogram of cryo-FIB milled MEFs. The lamella has a thickness of ∼130 nm. Missing wedge deconvolution of the tomogram was performed with IsoNet [46]. As a result, the platinum depositions on the surface of the lamella (yellow dashed rectangle) appear globular and VIF cross-sections (yellow arrow) reveal their pentameric shape. (**B**) More examples for the pentameric cross-sections of VIFs. Scale bar 10 nm.

**Extended Data Figure 2.**
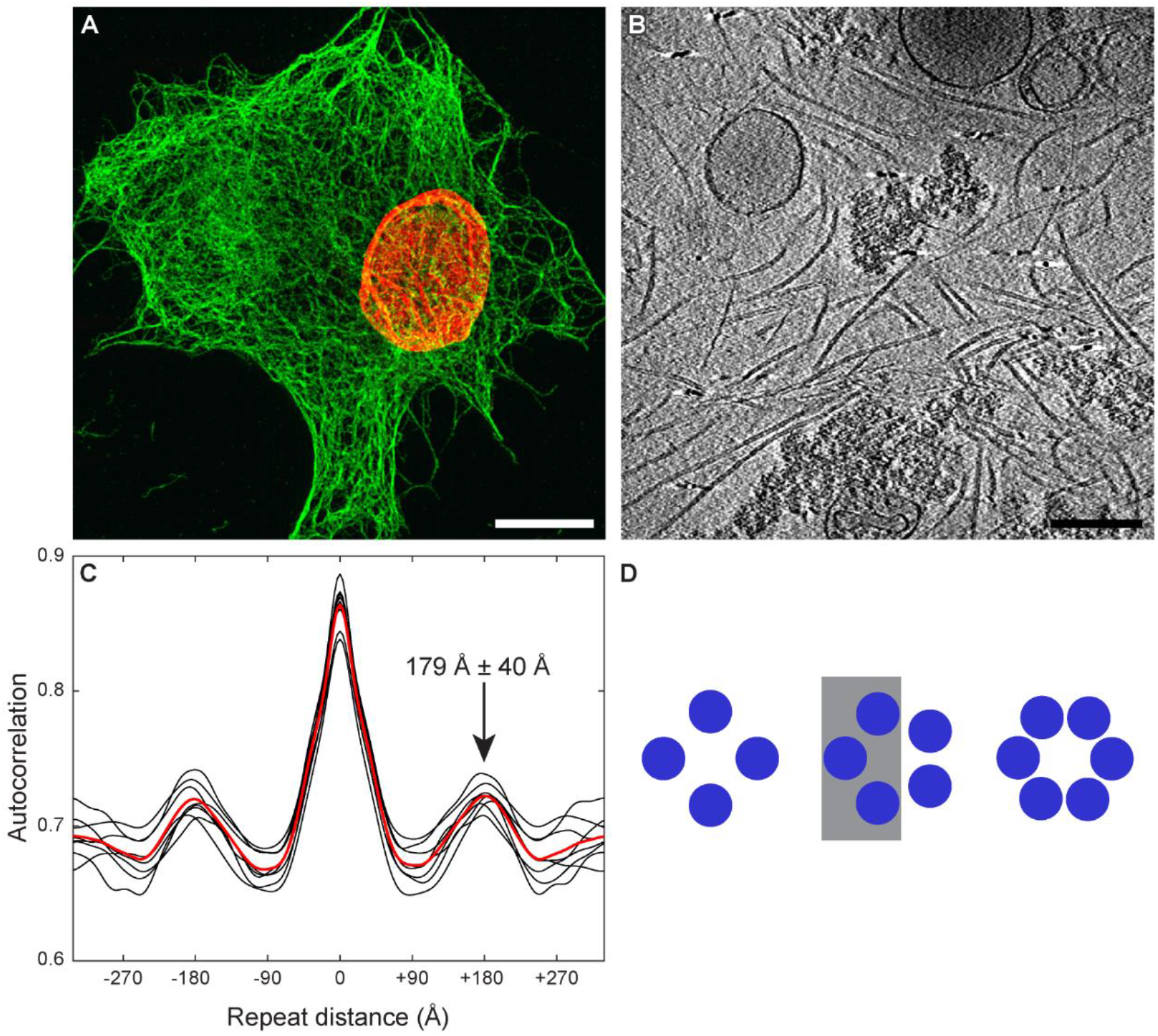
Cryo-ET of detergent-treated MEFs. (**A**) Maximum intensity projection of a 3D-SIM image of a detergent-treated MEF. The VIF network remains intact following the permeabilization procedure, fixation and staining with vimentin antibody (green). The cell nucleus is stained in red using lamin antibody. Scale bar 10 µm. (**B**) Slice in x-y-direction with 13.76 Å thickness through a cryo-tomogram of detergent-treated MEFs. Scale bar 200 nm. (**C**) The correlation of each class average with itself was calculated and the corresponding autocorrelation profiles along the x-axis were plotted (black lines). The red line is the averaged autocorrelation profile over all class averages. The pattern in the averages repeats at a distance of 179 Å ± 40 Å. (**D**) Model to explain the boundary asymmetry in the class averages. If VIFs are assembled from 5 protofibrils, one side of the VIF would appear brighter in projection (grey rectangle). However, if they are assembled from 4 or 6 protofibrils the VIF boundaries would have similar densities in projection.

**Extended Data Figure 3.**
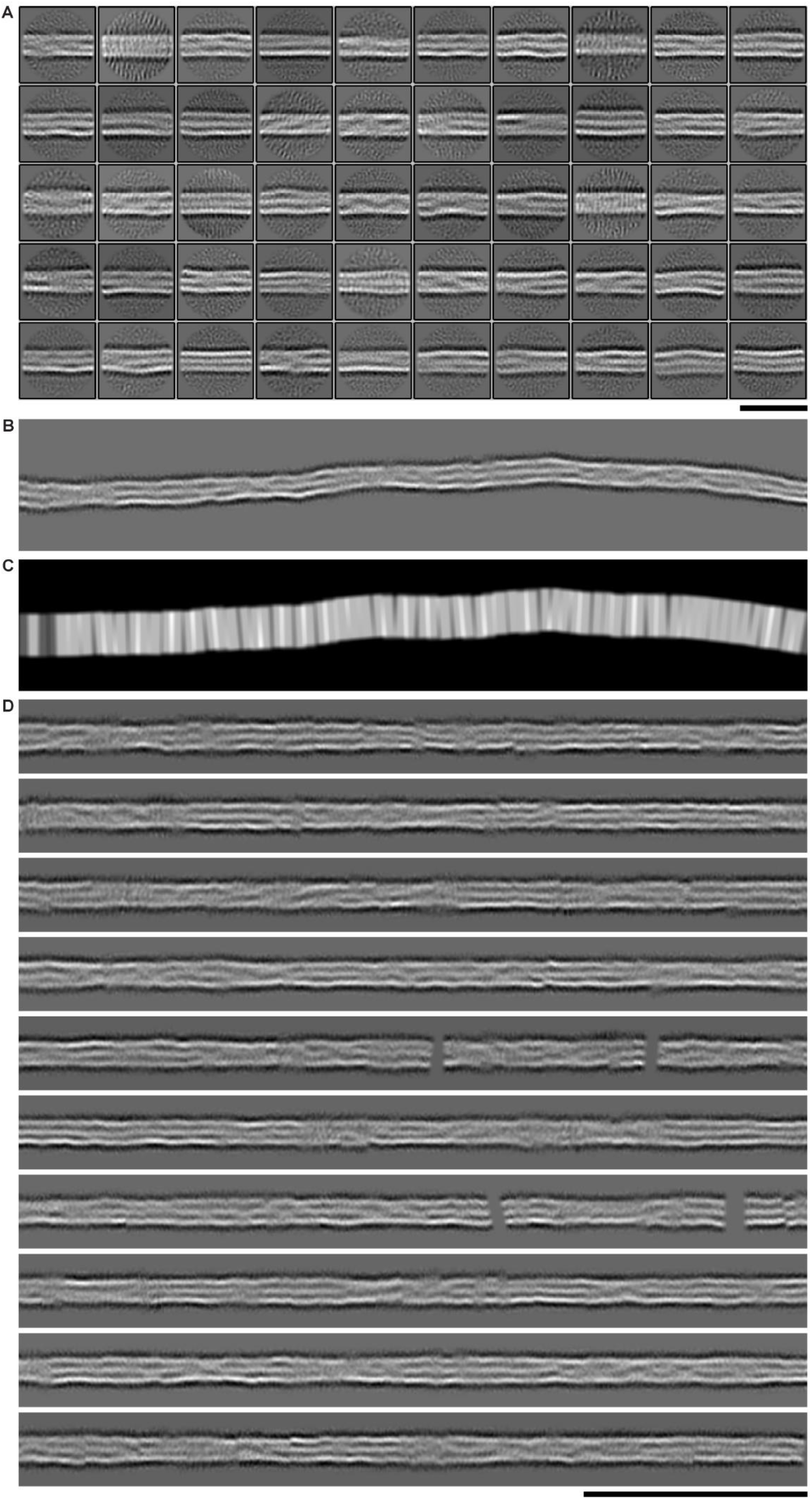
Computational assembly of VIFs. (**A**) VIF segments (615,106 particles with a size of 38 x 38 nm^2^) were extracted from tomograms of detergent-treated MEFs and combined with 2D classification. The picking distance between the segments was set to 55 Å and the projection thickness to 220 Å [48]. The displayed class averages were used for subsequent computational filament assembly. Scale bar 35 nm. (**B**) The computationally assembled VIFs (ca-VIFs) allow to follow the progression of the filaments with improved signal-to-noise ratio over a substantial length. The displayed filament box is 353 nm wide. (**C**) The ca-VIFs are represented by a series of transformed, tailored, and overlapping class averages. The densities are normalized according to the local overlap of the class averages. The image shows the normalization mask that was applied to the ca-VIF shown above. More overlap between the class averages is indicated by brighter regions. (**D**) Gallery of unbent ca-VIFs. Scale bar 100 nm.

**Extended Data Figure 4.**
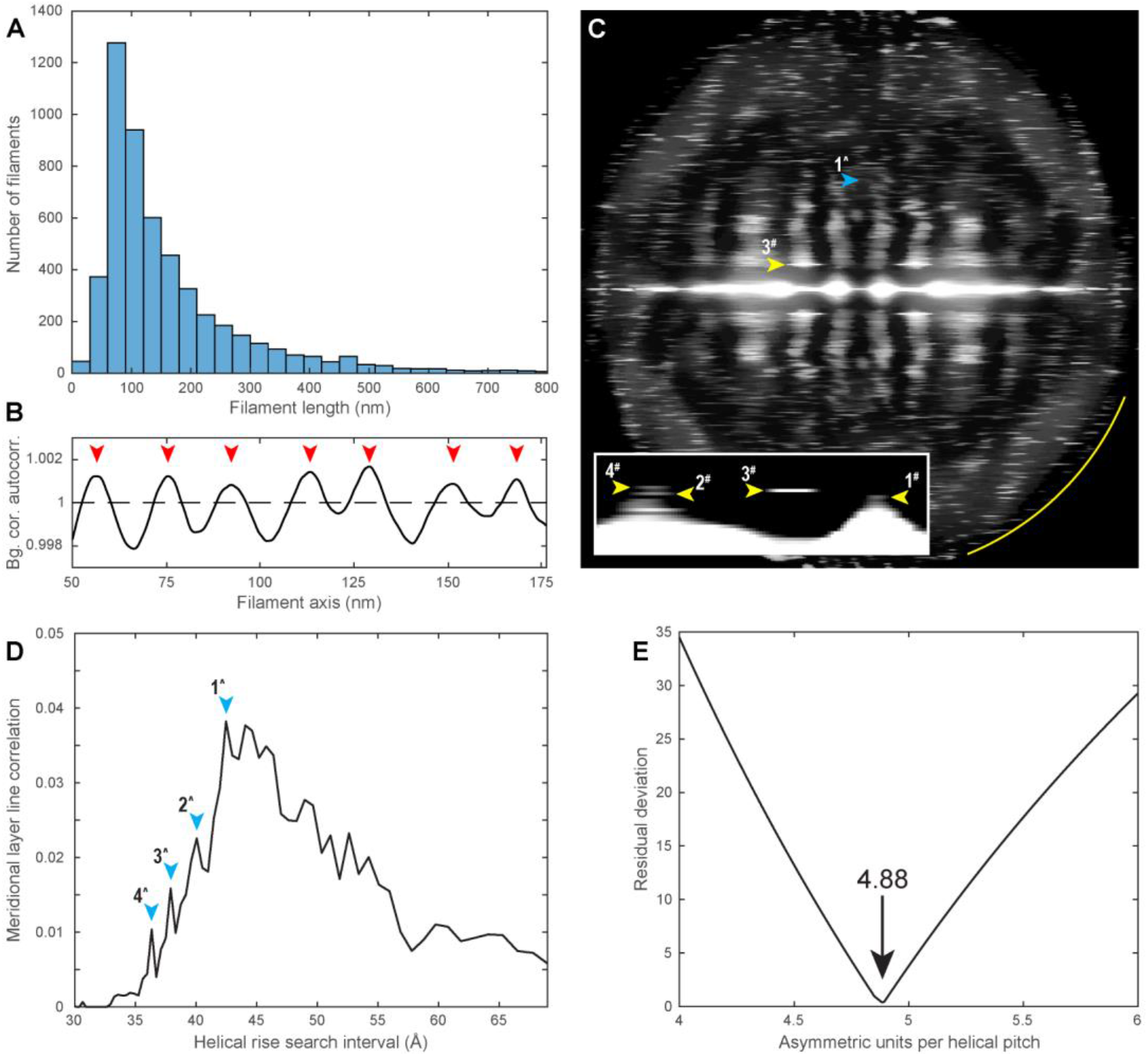
Helical parameter determination of VIFs. (**A**) Length histogram of the ca-VIFs. (**B**) Profile plot showing the averaged, background corrected autocorrelation signal of the ca-VIFs. The mean distance between the red arrowheads is 186.5 Å ± 26.0 Å. (**C**) In total, 389 ca-VIFs with a minimal length of 353°nm were combined into a power spectrum. The meridional reflection at 1/42.5 Å (blue arrowhead, 1^^^) indicates the helical rise of VIFs. Around the distinct layer line at 1/185.6 Å (yellow arrowhead, 3^#^), the following sequence of layer lines was extracted, indicated by 1^#^-4^#^ in the inset: 1/207.4 Å, 1/195.9 Å, 1/185.6 Å, 1/176.3 Å. The yellow arc indicates 1/16 Å. (**D**) The meridional reflection at 1/42.5 Å (blue arrowhead, 1^^^) was identified by cross-correlation of the layer lines with a zero order Bessel function within a search interval between 1/30 Å and 1/69 Å. In the proximity of the meridional reflection the following sequence of layer lines was extracted, indicated by 1^-4^ in the plot: 1/42.5 Å, 1/40.1 Å, 1/37.9 Å, 1/36.3 Å. (**E**) The optimal number of asymmetric units per helical pitch that relates both sequences of layer lines was determined.

**Extended Data Figure 5.**
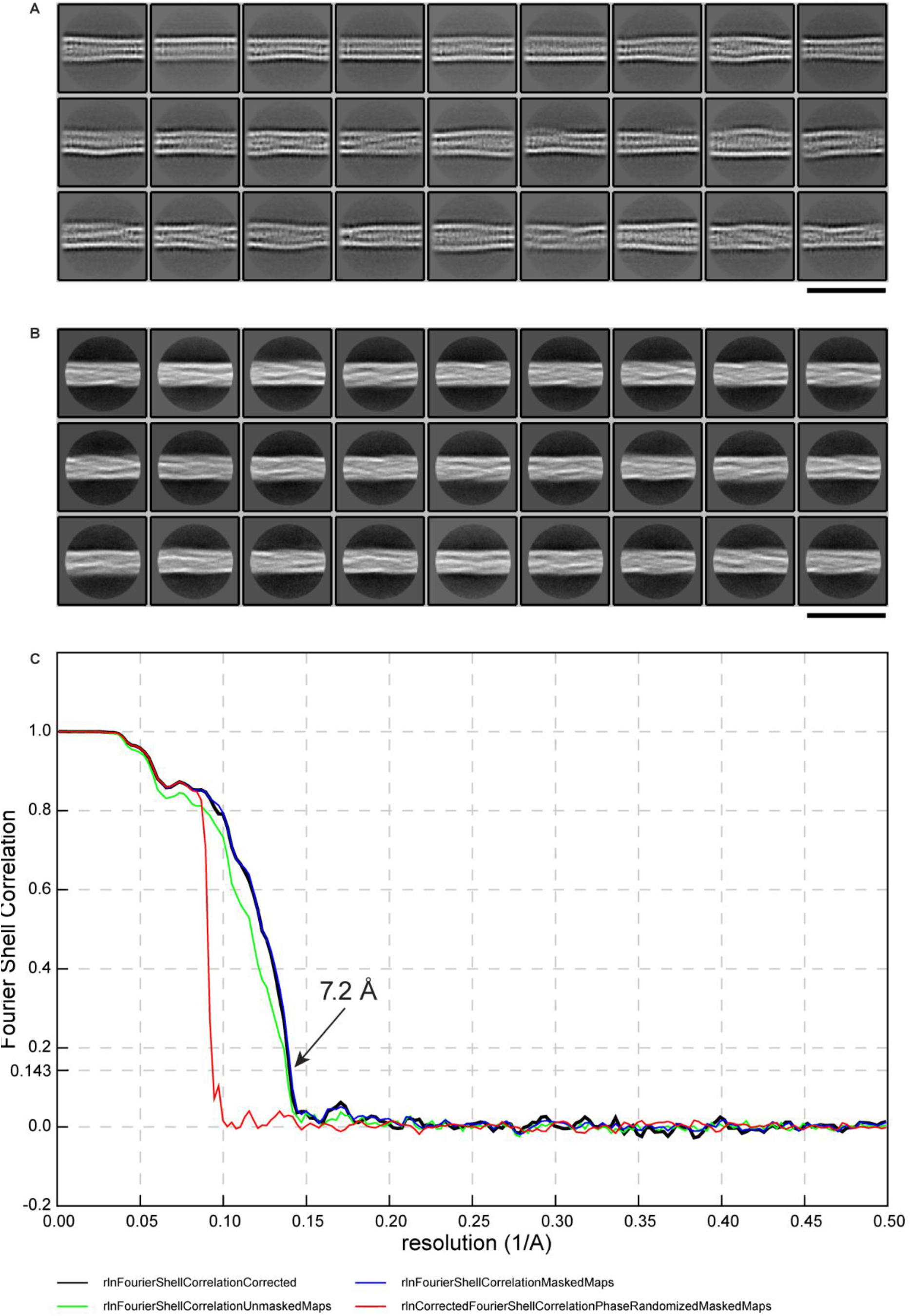
Cryo-EM data processing. (**A**) Before sorting the VIF segments with extensive 3D classifications, their 2D classes showed the previously detected pattern, namely that one filament wall appears more pronounced in projection than its counterpart. However, at this stage of data processing the VIF segments were considerably heterogenous regarding their diameter. The image shows a representative subset of 27 out of 200 class averages calculated from 801,585 VIF segments. Scale bar is 35 nm. (**B**) The VIF segments underlying the final VIF 3D structure were combined with 2D classification to assess the gain of structural homogeneity during data processing. The image shows a representative subset of 27 out of 100 class averages calculated from the final 236,920 VIF segments. Scale bar is 35 nm. (**C**) The gold-standard Fourier shell correlation (FSC) curve of the final VIF 3D structure indicates an overall resolution of 7.2 Å at FSC = 0.143.

**Extended Data Figure 6.**
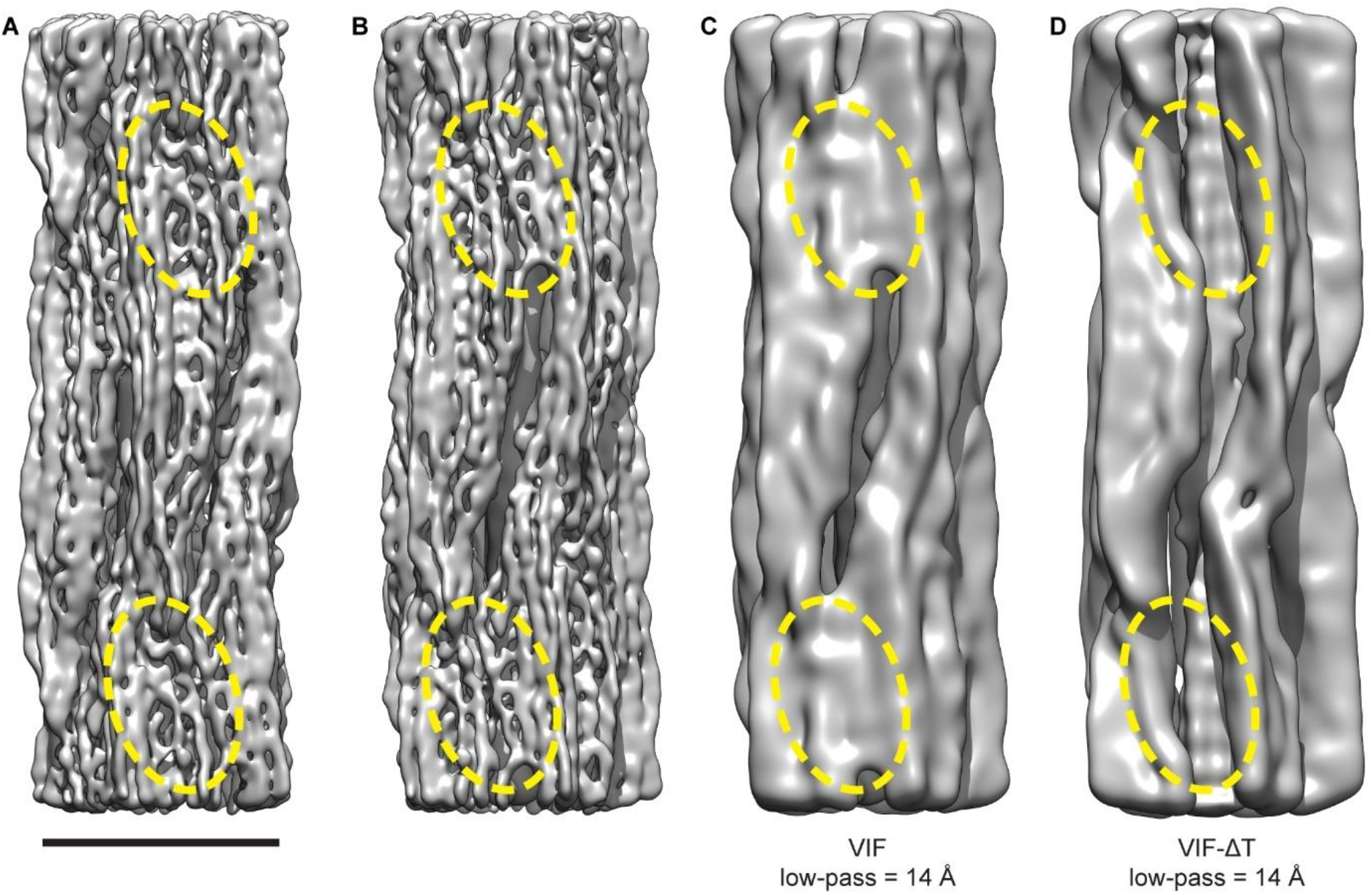
Comparison between the wildtype VIF and VIF-ΔT structures. (**A**) Isosurface rendering of the VIF structure. The contact sites between the protofibrils are circled by yellow dashed ovals. Scale bar is 10 nm. (**B**) Identical structure as shown in (**A**), but rotated -36° around the helical axis (**C**) Identical view as shown in (**B**), but the VIF structure was low-pass filtered to 14 Å. (**D**) Isosurface rendering of the VIF-ΔT structure, which is displayed with identical view and low-pass filter settings as the VIF structure depicted in (**C**). Both structures (VIF and VIF-ΔT) are similar to a great extent, and both structures contain a luminal fiber of similar volume. However, their striking difference, as indicated by the yellow dashed ovals in (**C**) and (**D**), is that the VIF-ΔT structure is clearly lacking the contact sites between the adjacent protofibrils.

**Extended Data Figure 7.**
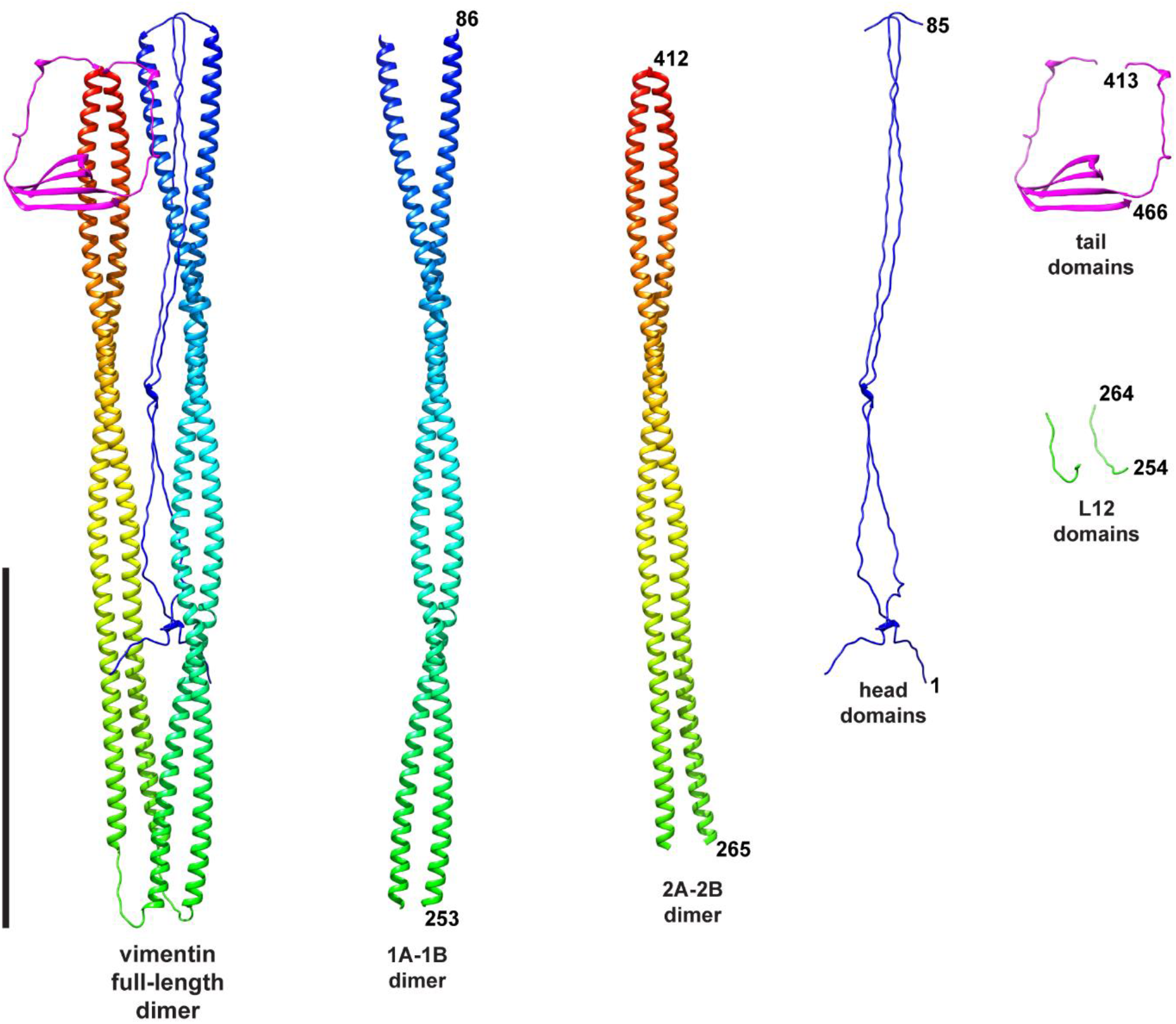
Alphafold prediction of the vimentin full-length dimer. The atomic model shown on the left is the ranked-0 alphafold prediction result of the vimentin full-length dimer. Based on this model we define the vimentin protein domains as follows: head, residues 1-85; 1A-1B, residues 86-253; L12, residues 254-264; 2A-2B, residues 265-412; tail, residues 413-466. Based on this definition the full-length dimer model was dissected into dimeric models of the vimentin protein domains (shown in the figure; labelled with 1A-1B dimer, 2A-2B dimer, head domains, tail domains, and L12 domains). These models constituted the initial folds for the subsequent building of the VIF atomic model. Scale bar 100 nm.

**Extended Data Figure 8.**
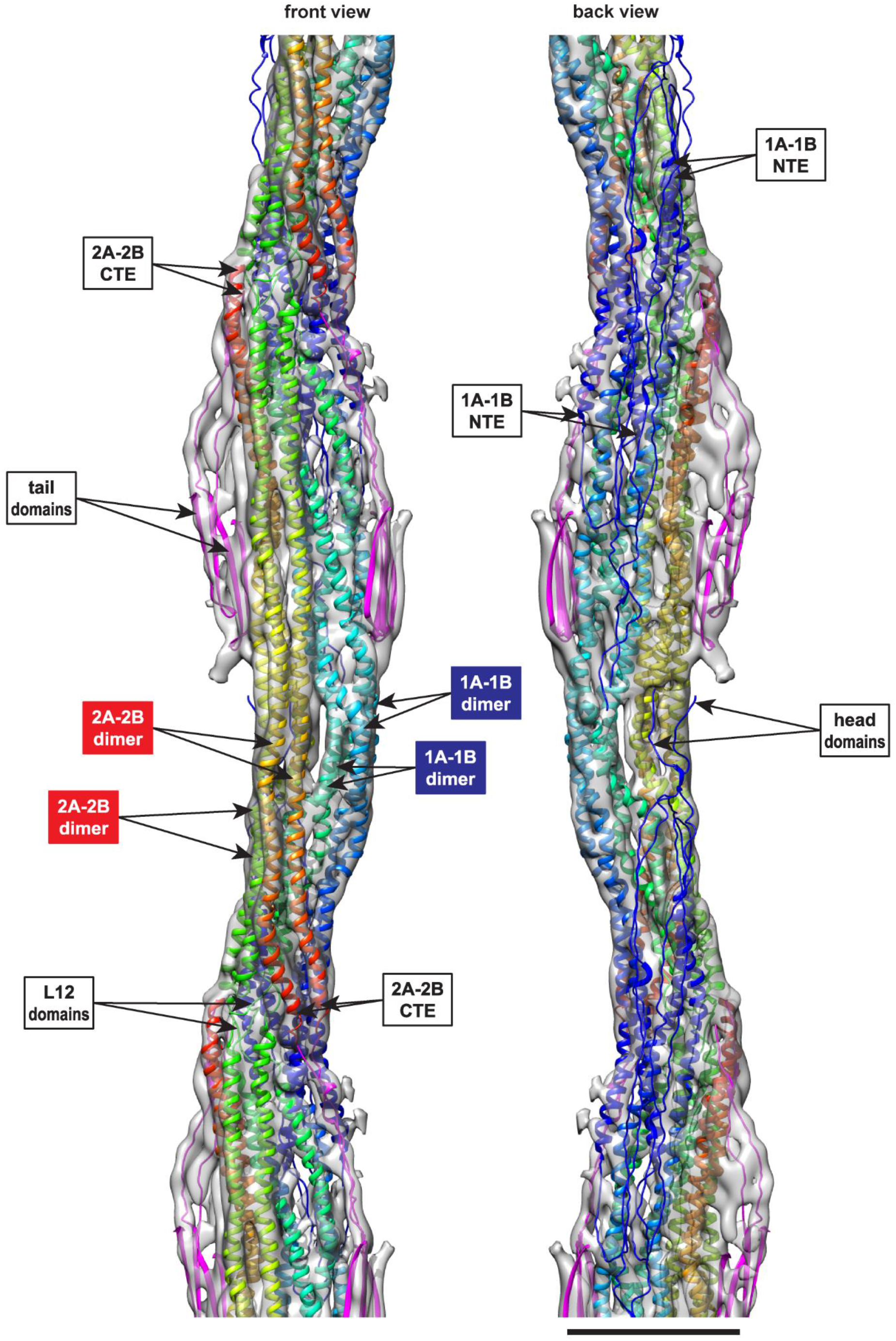
VIF 3D atomic model zoomed on protofibril. The complete VIF 3D atomic model was simplified to one repeating unit of a protofibril. The front view of this reduced model is displayed on the left, the view from the luminal face on the right, and the scale bar is 50 nm. The α-helical 2A-2B domains are colored from green**^NTE^**to red**^CTE^**, and the α-helical 1A-1B domains are colored from blue**^NTE^** to mint green**^CTE^**. The tail domains are colored magenta, the head domains blue, and the linker L12 domains green. The main constituents of the repeating unit of a protofibril are 2 antiparallel 2A-2B dimers (red labels) and 2 antiparallel 1A-1B dimers (blue labels). Along a protofibril, each of the 2A-2B dimers are connected by the L12 linker domains with one of the previous and one of the subsequent 1A-1B dimers. The CTEs of each 2A-2B dimer position the tail domains to form the lateral contact sites between the protofibrils, and the NTEs of each 1A-1B dimer position the head domains to form the luminal fiber.

**Extended Data Figure 9.**
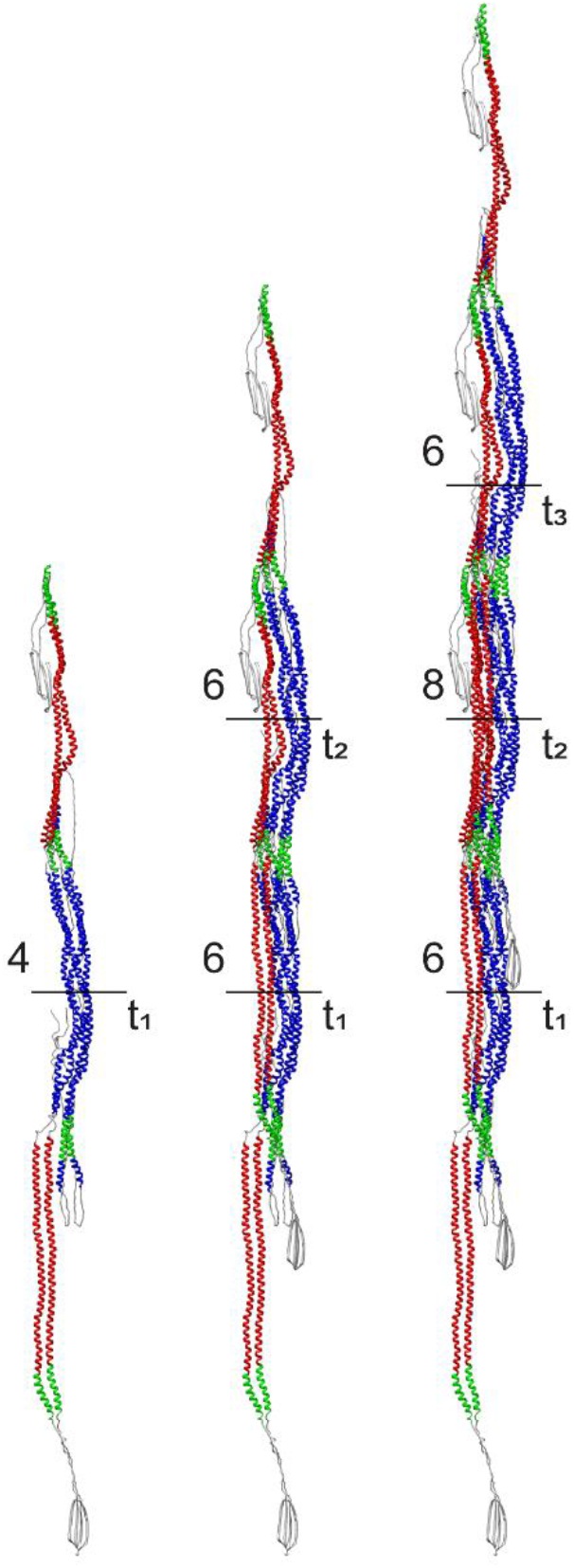
Construction of a protofibril. The number of polypeptide chains in cross-section is indicated by the numbers next to the horizontal black lines. Starting from a first tetramer t_1_ (shown on the left), a second tetramer t_2_ attaches with its 1A-1B section laterally to one of the flanking 2A-2B dimers of t_1_ (shown in the middle). This creates an intermediate assembly with 6 chains in cross-section. Subsequently, a third tetramer t_3_ binds to either side of this assembly (shown on the right). This creates a minimal length (∼110 nm) protofibril with its basic repeating unit (8 chains in cross-section) fully assembled in its center. Therefore, the full assembly of a protofibril with 8 polypeptide chains in cross-section requires the sequential interaction of at least 3 tetramers.

**Extended Data Figure 10.**
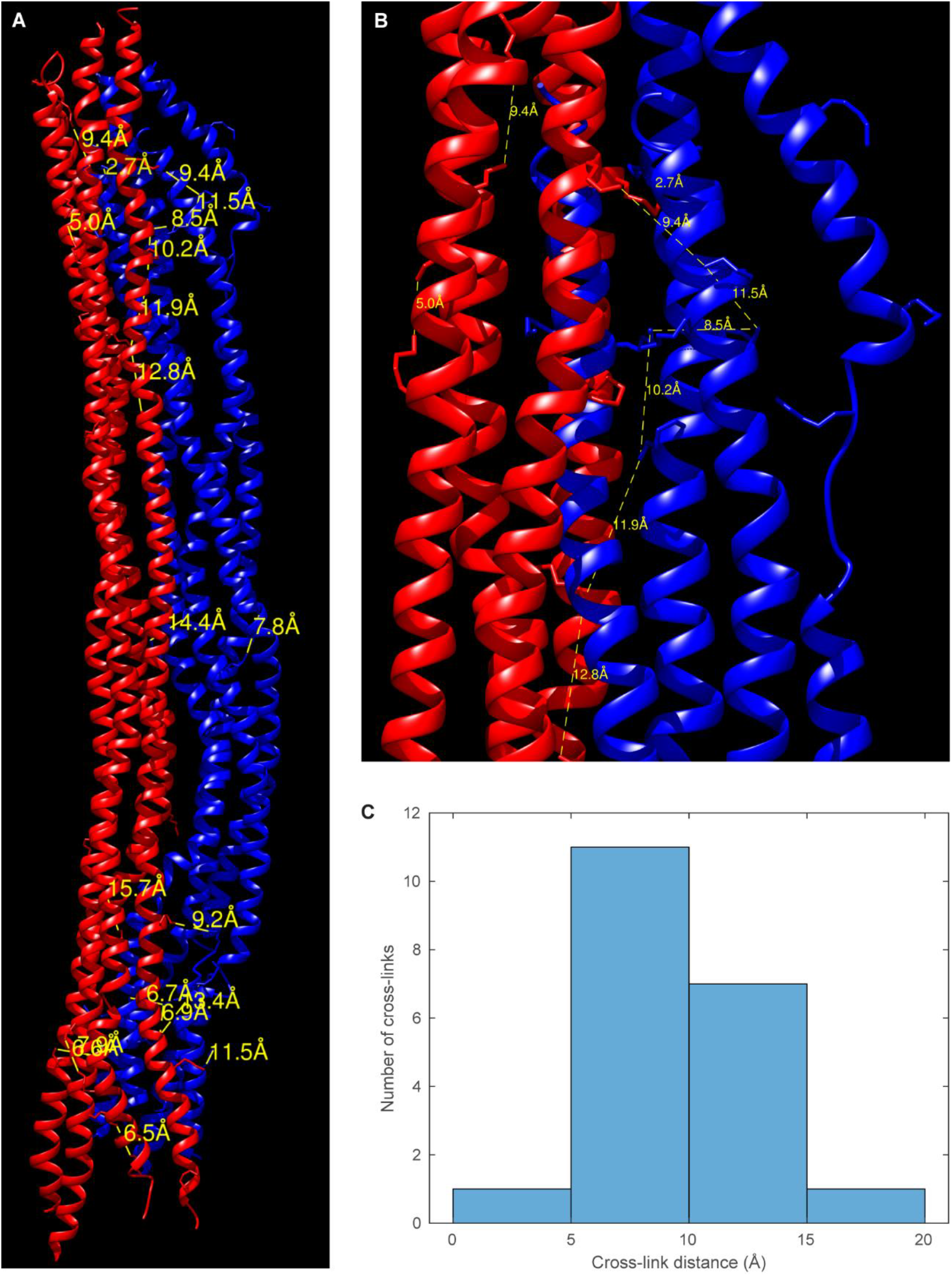
Mapping of cross-links. (**A**) Previously reported cross-links (yellow dashed lines) were superpositioned onto a model of the protofibril repeating unit (2A-2B dimers colored red, 1A-1B dimers colored blue). The cross-links were taken from references [61, 62]. (**B**) Zoom into the interlock region. (**C**) Histogram of the distances between the cross-linked residues in the model. The mean distance is 9.4 Å.

**Extended Data Figure 11.**
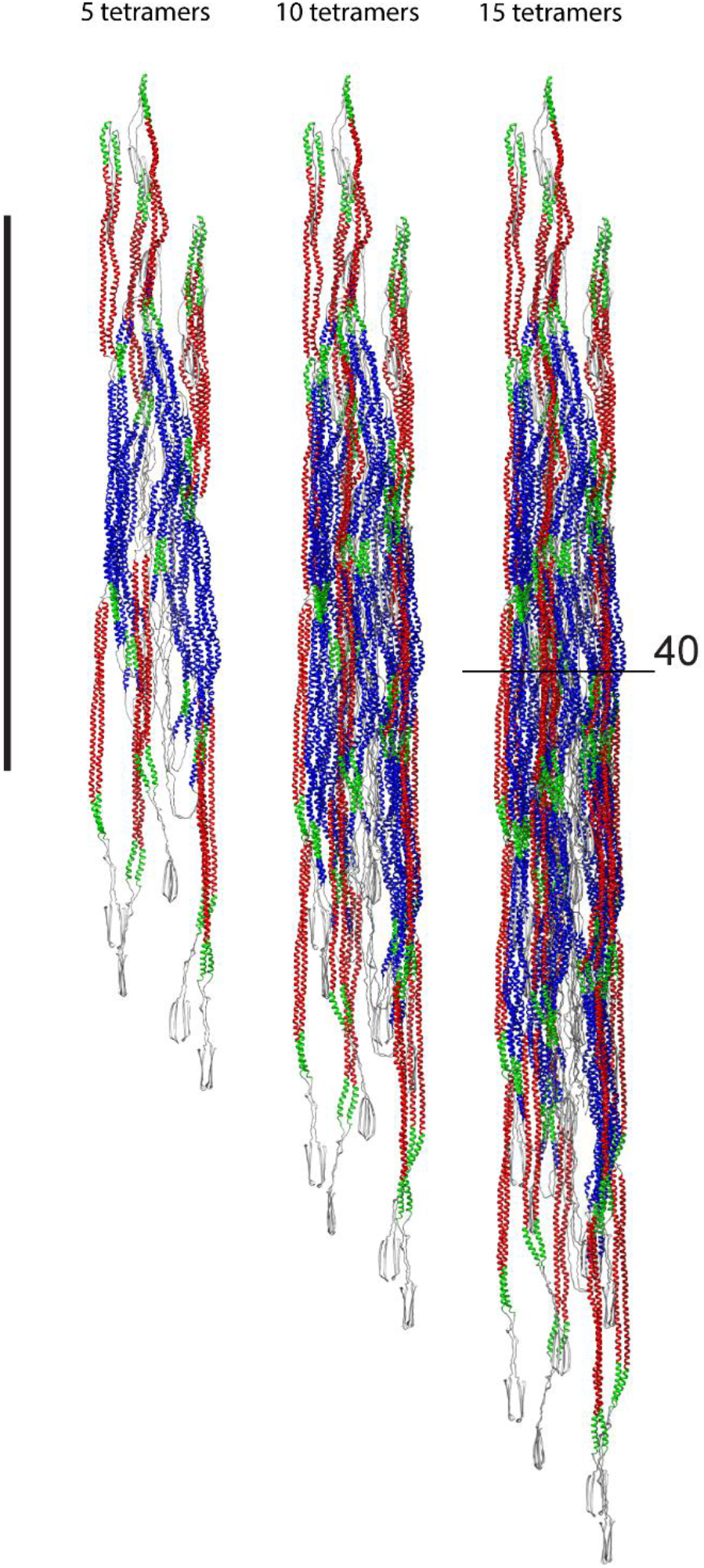
VIF models with increasing numbers of tetramers. (**A**) VIF model constructed from 5 tetramers. Scale bar is 50 nm. (**B**) VIF model constructed from 10 tetramers. (**C**) VIF model constructed from 15 tetramers. In-situ VIFs assembled from at least 15 tetramers exhibit 40 polypeptide chains in cross-section through their central regions.

**Extended Data Figure 12.**
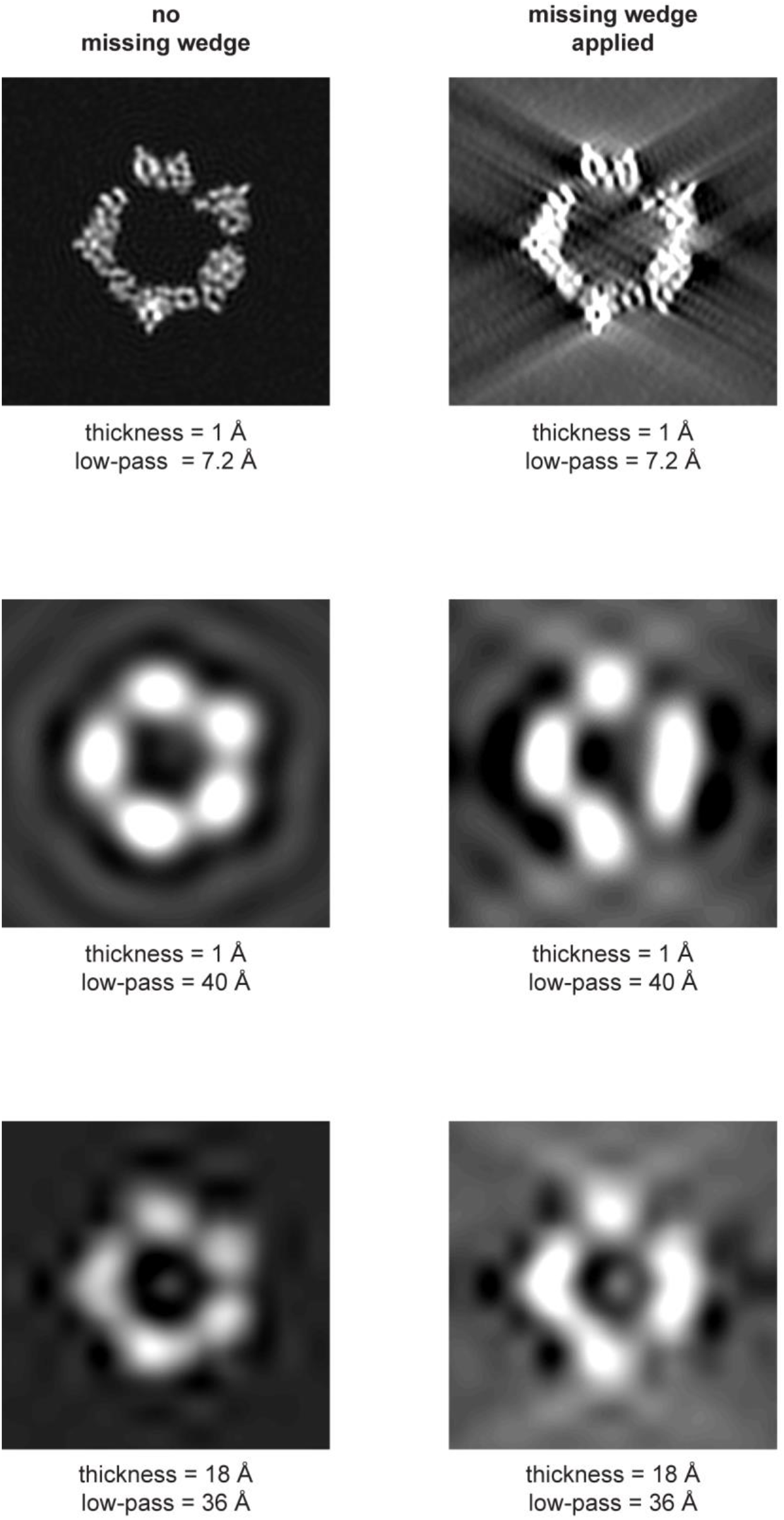
Missing wedge model calculation. The model calculation shows that cryo-ET without missing wedge deconvolution can resolve only 4 protofibrils in cross-section (right column). In the last row thickness and low-pass filter were chosen to reproduce the cross-sections displayed in reference [47].

## Extended Data Tables

**Extended Data Table 1.**
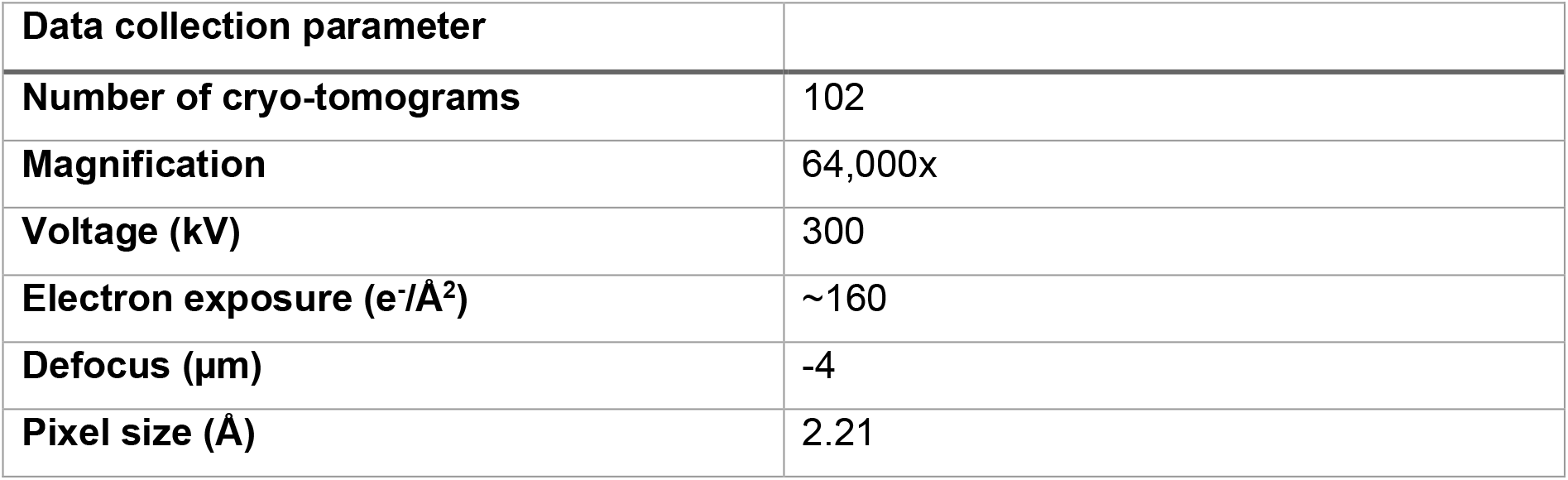
Cryo-FIB/Cryo-ET in-situ analysis of VIFs.

**Extended Data Table 2.** Cryo-ET of detergent-treated MEFs.

| **Data collection parameter** |  |
| --- | --- |
| **Number of cryo-tomograms** | 225 |
| **Magnification** | 42,000x |
| **Voltage (kV)** | 300 |
| **Electron exposure (e<sup>-</sup>/Å<sup>2</sup>)** | ~125 |
| **Defocus range (μm)** | -2 to -6 |
| **Pixel size (Å)** | 3.44 |
| **Class averages (65 x 65 nm<sup>2</sup>)** |  |
| **Initial particle images (no.)** | 390,297 |
| **Final particle images (no.)** | 133,780 |
| **Class averages (38 x 38 nm<sup>2</sup>)** |  |
| **Initial particle images (no.)** | 1,148,072 |
| **Final particle images (no.)** | 615,106 |
| **Computational filament assembly** |  |
| **Initial filaments (no.)** | 5,205 |
| **Final filaments (no.)** | 389 |

**Extended Data Table 3.**
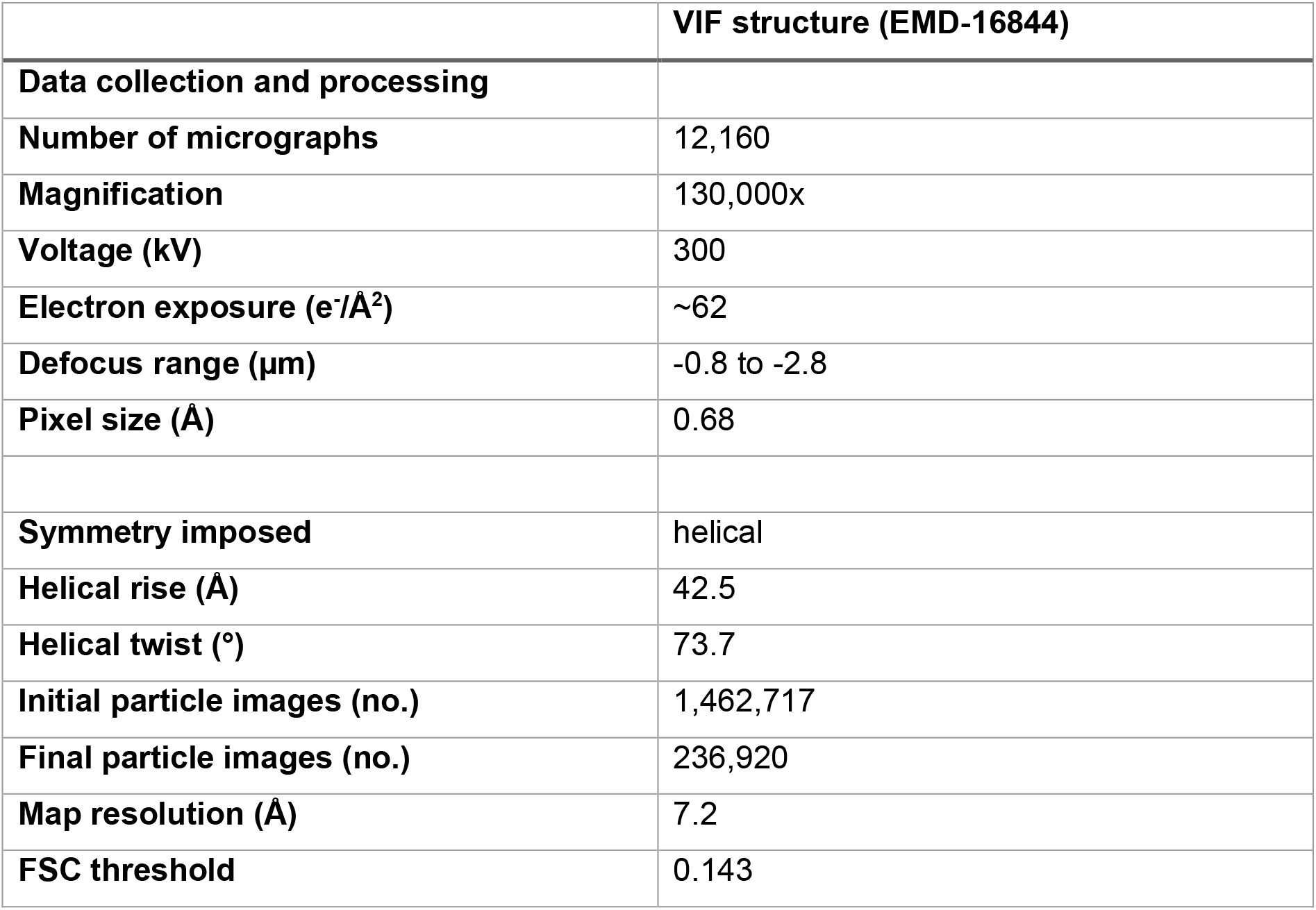
Cryo-EM of in-vitro polymerized human VIFs.

**Extended Data Table 4.** Cryo-EM of in-vitro polymerized human VIFs-ΔT.

|  | VIF-ΔT structure |
| --- | --- |
| **Data collection and processing** |  |
| **Number of micrographs** | 19,534 |
| **Magnification** | 130,000x |
| **Voltage (kV)** | 300 |
| **Electron exposure (e<sup>-</sup>/Å<sup>2</sup>)** | ~62 |
| **Defocus range (μm)** | -0.8 to -2.8 |
| **Pixel size (Å)** | 0.68 |
| **Symmetry imposed** | helical |
| **Helical rise (Å)** | 42.5 |
| **Helical twist (°)** | 73.7 |
| **Initial particle images (no.)** | 1,019,393 |
| **Final particle images (no.)** | 143,270 |
| **Map resolution (Å)** | 14 |
| **FSC threshold** | 0.143 |

## Supplementary Videos

**Supplementary Video 1. Tomogram of a cryo-FIB milled MEF.** The video shows successive slices of the IsoNet [46] corrected tomogram of a cryo-FIB milled MEF (**Fig. 1B**) from which the displayed VIF cross-sections were extracted (**Fig. 1D, Extended Data Fig. 1B**).

**Supplementary Video 2. Tomogram of a detergent-treated MEF.** The video shows successive slices of a tomogram of a detergent-treated MEF (**Extended Data Fig. 2B**).

**Supplementary Video 3. VIF structure.** The video shows the VIF structure (**Fig. 2A**) rotating around the helical axis.

**Supplementary Video 4. VIF atomic model.** The video shows the atomic model of VIFs (**Fig. 3A**) rotating around the helical axis. The 2A-2B dimers are colored from green**^NTE^** to red**^CTE^**. The 1A-1B dimers are colored from blue**^NTE^** to mint green**^CTE^**. The head domains are colored blue, the tail domains are colored magenta, the linker L12 domains are colored green.

**Supplementary Video 5. VIF atomic model cross-sections.** The video shows successive cross-sections of the VIF model (**Fig. 3D**). The movies ranges over a length of 207 Å.

**Supplementary Video 6. Extended VIF model.** The video shows a VIF model (**Fig. 4D**), assembled from 30 vimentin tetramers, rotating around the helical axis. The 2A-2B domains are colored red, the 1A-1B domains are colored blue, the between IFs highly conserved 1A and 1B domains are colored green.

## References

1. Steinert, P.M., A.C. Steven, and D.R. Roop, The molecular biology of intermediate filaments. Cell, 1985. 42(2): p. 411–20.

2. Szeverenyi, I., et al., The Human Intermediate Filament Database: comprehensive information on a gene family involved in many human diseases. Hum Mutat, 2008. 29(3): p. 351–60.

3. Turgay, Y., et al., The molecular architecture of lamins in somatic cells. Nature, 2017. 543(7644): p. 261-264.

4. Weber, M.S., et al., Structural heterogeneity of cellular K5/K14 filaments as revealed by cryo-electron microscopy. Elife, 2021. 10.

5. Khalil, M., et al., Neurofilaments as biomarkers in neurological disorders. Nature Reviews Neurology, 2018. 14(10): p. 577–589.

6. Herrmann, H., et al., Intermediate filaments: from cell architecture to nanomechanics. Nat Rev Mol Cell Biol, 2007. 8(7): p. 562–73.

7. Sanghvi-Shah, R. and G.F. Weber, Intermediate Filaments at the Junction of Mechanotransduction, Migration, and Development. Frontiers in Cell and Developmental Biology, 2017. 5.

8. Omary, M.B., P.A. Coulombe, and W.H.I. McLean, Mechanisms of disease: Intermediate filament proteins and their associated diseases. New England Journal of Medicine, 2004. 351(20): p. 2087–2100.

9. Vikstrom, K.L., et al., Steady state dynamics of intermediate filament networks. J Cell Biol, 1992. 118(1): p. 121–9.

10. Hu, J., et al., High stretchability, strength, and toughness of living cells enabled by hyperelastic vimentin intermediate filaments. Proc Natl Acad Sci U S A, 2019. 116(35): p. 17175–17180.

11. Kreplak, L., et al., Exploring the mechanical behavior of single intermediate filaments. J Mol Biol, 2005. 354(3): p. 569–77.

12. Block, J., et al., Nonlinear Loading-Rate-Dependent Force Response of Individual Vimentin Intermediate Filaments to Applied Strain. Phys Rev Lett, 2017. 118(4): p. 048101.

13. Janmey, P.A., et al., Viscoelastic properties of vimentin compared with other filamentous biopolymer networks. J Cell Biol, 1991. 113(1): p. 155–60.

14. Goldman, R.D., et al., The function of intermediate filaments in cell shape and cytoskeletal integrity. J Cell Biol, 1996. 134(4): p. 971–83.

15. Lowery, J., et al., Intermediate Filaments Play a Pivotal Role in Regulating Cell Architecture and Function. J Biol Chem, 2015. 290(28): p. 17145–53.

16. Tsuruta, D. and J.C. Jones, The vimentin cytoskeleton regulates focal contact size and adhesion of endothelial cells subjected to shear stress. J Cell Sci, 2003. 116(Pt 24): p. 4977–84.

17. Helfand, B.T., et al., Vimentin organization modulates the formation of lamellipodia. Mol Biol Cell, 2011. 22(8): p. 1274–89.

18. Jiu, Y., et al., Vimentin intermediate filaments control actin stress fiber assembly through GEF-H1 and RhoA. J Cell Sci, 2017. 130(5): p. 892–902.

19. Wu, H.Y., et al., Vimentin intermediate filaments and filamentous actin form unexpected interpenetrating networks that redefine the cell cortex. Proceedings of the National Academy of Sciences of the United States of America, 2022. 119(10).

20. Perlson, E., et al., Vimentin-dependent spatial translocation of an activated MAP kinase in injured nerve. Neuron, 2005. 45(5): p. 715–726.

21. Mendez, M.G., S. Kojima, and R.D. Goldman, Vimentin induces changes in cell shape, motility, and adhesion during the epithelial to mesenchymal transition. FASEB J, 2010. 24(6): p. 1838–51.

22. Kidd, M.E., D.K. Shumaker, and K.M. Ridge, The role of vimentin intermediate filaments in the progression of lung cancer. Am J Respir Cell Mol Biol, 2014. 50(1): p. 1–6.

23. Satelli, A. and S.L. Li, Vimentin in cancer and its potential as a molecular target for cancer therapy. Cellular and Molecular Life Sciences, 2011. 68(18): p. 3033–3046.

24. Sivagurunathan, S., et al., Expression of vimentin alters cell mechanics, cell-cell adhesion, and gene expression profiles suggesting the induction of a hybrid EMT in human mammary epithelial cells. Frontiers in Cell and Developmental Biology, 2022. 10.

25. Danielsson, F., et al., Vimentin Diversity in Health and Disease. Cells, 2018. 7(10).

26. Muller, M., et al., Dominant cataract formation in association with a vimentin assembly disrupting mutation. Hum Mol Genet, 2009. 18(6): p. 1052–7.

27. Henderson, P., et al., A role for vimentin in Crohn disease. Autophagy, 2012. 8(11): p. 1695–6.

28. Fernandez-Ortega, C., et al., Identification of Vimentin as a Potential Therapeutic Target against HIV Infection. Viruses-Basel, 2016. 8(6).

29. Strelkov, S.V., et al., Conserved segments 1A and 2B of the intermediate filament dimer: their atomic structures and role in filament assembly. EMBO J, 2002. 21(6): p. 1255–66.

30. Parry, D.A. and P.M. Steinert, Intermediate filaments: molecular architecture, assembly, dynamics and polymorphism. Q Rev Biophys, 1999. 32(2): p. 99–187.

31. Lilina, A.V., et al., Stability profile of vimentin rod domain. Protein Sci, 2022. 31(12): p. e4505.

32. Sokolova, A.V., et al., Monitoring intermediate filament assembly by small-angle x-ray scattering reveals the molecular architecture of assembly intermediates. Proc Natl Acad Sci U S A, 2006. 103(44): p. 16206–11.

33. Chernyatina, A.A., et al., Atomic structure of the vimentin central alpha-helical domain and its implications for intermediate filament assembly. Proc Natl Acad Sci U S A, 2012. 109(34): p. 13620–5.

34. Qin, Z., L. Kreplak, and M.J. Buehler, Hierarchical structure controls nanomechanical properties of vimentin intermediate filaments. PLoS One, 2009. 4(10): p. e7294.

35. Kirmse, R., et al., A quantitative kinetic model for the in vitro assembly of intermediate filaments from tetrameric vimentin. J Biol Chem, 2007. 282(25): p. 18563–18572.

36. Noding, B., H. Herrmann, and S. Koster, Direct observation of subunit exchange along mature vimentin intermediate filaments. Biophys J, 2014. 107(12): p. 2923–2931.

37. Robert, A., et al., Vimentin filament precursors exchange subunits in an ATP-dependent manner. Proc Natl Acad Sci U S A, 2015. 112(27): p. E3505–14.

38. Herrmann, H., et al., Structure and assembly properties of the intermediate filament protein vimentin: the role of its head, rod and tail domains. J Mol Biol, 1996. 264(5): p. 933–53.

39. Herrmann, H., et al., Characterization of distinct early assembly units of different intermediate filament proteins. J Mol Biol, 1999. 286(5): p. 1403–20.

40. Mucke, N., et al., Assembly Kinetics of Vimentin Tetramers to Unit-Length Filaments: A Stopped-Flow Study. Biophys J, 2018. 114(10): p. 2408–2418.

41. Eldirany, S.A., et al., Recent insight into intermediate filament structure. Current Opinion in Cell Biology, 2021. 68: p. 132–143.

42. Rigort, A., et al., Micromachining tools and correlative approaches for cellular cryo-electron tomography. J Struct Biol, 2010. 172(2): p. 169–79.

43. Rigort, A., et al., Focused ion beam micromachining of eukaryotic cells for cryoelectron tomography. Proc Natl Acad Sci U S A, 2012. 109(12): p. 4449–54.

44. Mahamid, J., et al., Visualizing the molecular sociology at the HeLa cell nuclear periphery. Science, 2016. 351(6276): p. 969-972.

45. Beck, M. and W. Baumeister, Cryo-Electron Tomography: Can it Reveal the Molecular Sociology of Cells in Atomic Detail? Trends Cell Biol, 2016. 26(11): p. 825–837.

46. Liu, Y.T., et al., Isotropic reconstruction for electron tomography with deep learning. Nat Commun, 2022. 13(1): p. 6482.

47. Goldie, K.N., et al., Dissecting the 3-D structure of vimentin intermediate filaments by cryo-electron tomography. J Struct Biol, 2007. 158(3): p. 378–85.

48. Martins, B., et al., Unveiling the polarity of actin filaments by cryo-electron tomography. Structure, 2021.

49. Bharat, T.A.M. and S.H.W. Scheres, Resolving macromolecular structures from electron cryo-tomography data using subtomogram averaging in RELION. Nature Protocols, 2016. 11(11): p. 9–20.

50. Diaz, R., W.J. Rice, and D.L. Stokes, Fourier-Bessel reconstruction of helical assemblies. Methods Enzymol, 2010. 482: p. 131–65.

51. Henderson, D., N. Geisler, and K. Weber, A periodic ultrastructure in intermediate filaments. J Mol Biol, 1982. 155(2): p. 173–6.

52. Milam, L. and H.P. Erickson, Visualization of a 21-nm axial periodicity in shadowed keratin filaments and neurofilaments. J Cell Biol, 1982. 94(3): p. 592–6.

53. Herrmann, H., et al., The intermediate filament protein consensus motif of helix 2B: its atomic structure and contribution to assembly. J Mol Biol, 2000. 298(5): p. 817–32.

54. Davies, D.B., et al., Structural molecular biology : methods and applications. NATO advanced study institutes series Series A, Life sciences. 1982, New York: Plenum Press. x, 530 p.

55. Scheres, S.H., RELION: implementation of a Bayesian approach to cryo-EM structure determination. J Struct Biol, 2012. 180(3): p. 519–30.

56. He, S. and S.H.W. Scheres, Helical reconstruction in RELION. J Struct Biol, 2017. 198(3): p. 163–176.

57. Kato, M., X. Zhou, and S.L. McKnight, How do protein domains of low sequence complexity work? RNA, 2022. 28(1): p. 3–15.

58. Jumper, J., et al., Highly accurate protein structure prediction with AlphaFold. Nature, 2021. 596(7873): p. 583-589.

59. Trabuco, L.G., et al., Flexible fitting of atomic structures into electron microscopy maps using molecular dynamics. Structure, 2008. 16(5): p. 673–83.

60. Kidmose, R.T., et al., Namdinator - automatic molecular dynamics flexible fitting of structural models into cryo-EM and crystallography experimental maps. IUCrJ, 2019. 6(Pt 4): p. 526–531.

61. Steinert, P.M., L.N. Marekov, and D.A. Parry, Diversity of intermediate filament structure. Evidence that the alignment of coiled-coil molecules in vimentin is different from that in keratin intermediate filaments. J Biol Chem, 1993. 268(33): p. 24916–25.

62. Parry, D.A., L.N. Marekov, and P.M. Steinert, Subfilamentous protofibril structures in fibrous proteins: cross-linking evidence for protofibrils in intermediate filaments. J Biol Chem, 2001. 276(42): p. 39253–8.

63. Rappsilber, J., The beginning of a beautiful friendship: cross-linking/mass spectrometry and modelling of proteins and multi-protein complexes. J Struct Biol, 2011. 173(3): p. 530–40.

64. Lin, Y., et al., Toxic PR Poly-Dipeptides Encoded by the C9orf72 Repeat Expansion Target LC Domain Polymers. Cell, 2016. 167(3): p. 789–802 e12.

65. Zhou, X.M., et al., Transiently structured head domains control intermediate filament assembly. Proceedings of the National Academy of Sciences of the United States of America, 2021. 118(8).

66. Meier, M., et al., Vimentin coil 1A-A molecular switch involved in the initiation of filament elongation. J Mol Biol, 2009. 390(2): p. 245–61.

67. Mastronarde, D.N., Automated electron microscope tomography using robust prediction of specimen movements. Journal of Structural Biology, 2005. 152(1): p. 36–51.

68. Hagen, W.J.H., W. Wan, and J.A.G. Briggs, Implementation of a cryo-electron tomography tilt-scheme optimized for high resolution subtomogram averaging. J Struct Biol, 2017. 197(2): p. 191–198.

69. Nickell, S., et al., TOM software toolbox: acquisition and analysis for electron tomography. J Struct Biol, 2005. 149(3): p. 227–34.

70. Li, X.M., et al., Electron counting and beam-induced motion correction enable near-atomic-resolution single-particle cryo-EM. Nature Methods, 2013. 10(6): p. 584-+.

71. Eibauer, M., et al., Unraveling the structure of membrane proteins in situ by transfer function corrected cryo-electron tomography. Journal of Structural Biology, 2012. 180(3): p. 488–496.

72. Chen, M., et al., Convolutional neural networks for automated annotation of cellular cryo-electron tomograms. Nat Methods, 2017. 14(10): p. 983–985.

73. Pettersen, E.F., et al., UCSF chimera - A visualization system for exploratory research and analysis. Journal of Computational Chemistry, 2004. 25(13): p. 1605–1612.

74. Kronenberg-Tenga, R., et al., A lamin A/C variant causing striated muscle disease provides insights into filament organization. Journal of Cell Science, 2021. 134(6): p. jcs256156.

75. Schroeder, A.B., et al., The ImageJ ecosystem: Open-source software for image visualization, processing, and analysis. Protein Sci, 2021. 30(1): p. 234–249.

76. Kocsis, E., et al., Image Averaging of Flexible Fibrous Macromolecules - the Clathrin Triskelion Has an Elastic Proximal Segment. Journal of Structural Biology, 1991. 107(1): p. 6–14.

77. Herrmann, H., L. Kreplak, and U. Aebi, Isolation, characterization, and in vitro assembly of intermediate filaments. Intermediate Filament Cytoskeleton, 2004. 78: p. 3-24.

78. Zhang, K., Gctf: Real-time CTF determination and correction. Journal of Structural Biology, 2016. 193(1): p. 1–12.

79. Wagner, T., et al., SPHIRE-crYOLO is a fast and accurate fully automated particle picker for cryo-EM. Communications Biology, 2019. 2.

80. Ramirez-Aportela, E., et al., Automatic local resolution-based sharpening of cryo-EM maps. Bioinformatics, 2020. 36(3): p. 765–772.

81. Kucukelbir, A., F.J. Sigworth, and H.D. Tagare, Quantifying the local resolution of cryo-EM density maps. Nat Methods, 2014. 11(1): p. 63–5.

82. Zheng, S.Q., et al., MotionCor2: anisotropic correction of beam-induced motion for improved cryo-electron microscopy. Nat Methods, 2017. 14(4): p. 331–332.

83. Kozakov, D., et al., The ClusPro web server for protein-protein docking. Nat Protoc, 2017. 12(2): p. 255–278.

84. Trabuco, L.G., et al., Molecular dynamics flexible fitting: a practical guide to combine cryo-electron microscopy and X-ray crystallography. Methods, 2009. 49(2): p. 174–80.

85. Pintilie, G. and W. Chiu, Comparison of Segger and other methods for segmentation and rigid-body docking of molecular components in cryo-EM density maps. Biopolymers, 2012. 97(9): p. 742–60.

86. Oosterheert, W., et al., Structural basis of actin filament assembly and aging. Nature, 2022. 611(7935): p. 374-379.

87. Zhang, R., B. LaFrance, and E. Nogales, Separating the effects of nucleotide and EB binding on microtubule structure. Proc Natl Acad Sci U S A, 2018. 115(27): p. E6191–E6200.

